# Intra-ripple frequency accommodation in an inhibitory network model for hippocampal ripple oscillations

**DOI:** 10.1101/2023.01.30.526209

**Authors:** Natalie Schieferstein, Tilo Schwalger, Benjamin Lindner, Richard Kempter

**Affiliations:** Humboldt-Universität zu Berlin, Department of Biology, Institute for Theoretical Biology, 10115 Berlin, Germany; Bernstein Center for Computational Neuroscience Berlin, 10115 Berlin, Germany; Technische Universität Berlin, Institute for Mathematics, 10587 Berlin, Germany; Humboldt-Universität zu Berlin, Department of Physics, 12489 Berlin, Germany; Einstein Center for Neurosciences Berlin, 10117 Berlin, Germany

## Abstract

Hippocampal ripple oscillations have been implicated in important cognitive functions such as memory consolidation and planning. Multiple computational models have been proposed to explain the emergence of ripple oscillations, relying either on excitation or inhibition as the main pacemaker. Nevertheless, the generating mechanism of ripples remains unclear. An interesting dynamical feature of experimentally measured ripples, which may advance model selection, is intra-ripple frequency accommodation (IFA): a decay of the instantaneous ripple frequency over the course of a ripple event. So far, only the “bifurcation-based” inhibition-first model, which is based on delayed inhibitory synaptic coupling, has been shown to reproduce IFA. Here we use an analytical mean-field approach and numerical simulations of a leaky integrate-and-fire spiking network to explain the mechanism of IFA. We develop a drift-based approximation for the oscillation dynamics of the population rate and the mean membrane potential of interneurons under strong drive and strong inhibitory coupling. We demonstrate that IFA arises due to a speed-dependent hysteresis effect in the dynamics of the mean membrane potential, when the interneurons receive transient, sharp wave-associated excitation. We thus predict that the IFA asymmetry vanishes in the limit of slowly changing drive, but is otherwise a robust feature of the bifurcation-based inhibition-first ripple model.

## Introduction

The brain exhibits a variety of oscillations across brain areas and states (Berger, 1933; Buzsáki, 2006; Steriade et al., 1987, 1993; Vanderwolf, 1969; Buzsáki et al., 1983; Colgin, 2016; O’Keefe, 1976; Buzsáki et al., 1992). Different rhythms have been shown to correlate with different cognitive functions such as the encoding of new information (Colgin, 2016), communication between brain areas (Sirota et al., 2008; Oyanedel et al., 2020; Fries, 2005, but see Schneider et al., 2021) or the consolidation of memories (Girardeau et al., 2009; Ego-Stengel and Wilson, 2010). It is therefore important to understand the neuronal mechanisms behind the generation of oscillations.

Hippocampal sharp wave-ripples (SPW-R) in particular have been associated with the replay of behaviorally relevant neuronal activity (Lee and Wilson, 2002; Diba and Buzsáki, 2007; Vaz et al., 2020) and might hence be important for memory consolidation (Buzsáki, 1989; Nádasdy et al., 1999; Lee and Wilson, 2002; Girardeau et al., 2009; Ego-Stengel and Wilson, 2010; Fernández-Ruiz et al., 2019) and planning (Diba and Buzsáki, 2007; Carr et al., 2011; Jadhav et al., 2012; Pfeiffer and Foster, 2013).

SPW-Rs are transient events (duration 50–100 ms, incidence ∼ 1/second) that can be measured in the local field potential (LFP) of the hippocampus *in vivo* during slow wave sleep or quiet rest (O’Keefe, 1976; Buzsáki et al., 1992) as well as *in vitro* (Wu et al., 2002; Maier et al., 2003; Maier and Kempter, 2017). A SPW-R event is marked by a transient increase in spiking activity (reflected in the sharp wave) modulated by a fast oscillation (the ripple, 150–250 Hz *in vivo*, Buzsáki et al. (1992); 210 ± 16 Hz *in vitro*, Maier et al. (2003)). In CA1 the LFP sharp wave component is most prominent in layer *stratum radiatum* and is thought to reflect a current sink due to elevated excitatory synaptic transmission from the CA3 Schaffer collaterals. The ripple component is strongest in *stratum pyramidale* of CA1 and is thought to reflect inhibitory synaptic currents, and potentially rhythmic excitatory action potentials (Mitzdorf, 1985; Schomburg et al., 2012). Ripple oscillations can also be found in the upstream field CA3, but there they are generally weaker and slightly slower (Sullivan et al., 2011). Furthermore, slice preparations have shown that the isolated CA1 can generate ripple oscillations without oscillatory input from CA3 (Maier et al., 2003, 2011). It is therefore believed that the local CA1 network can generate ripples in isolation.

Various computational models have been put forward to explain ripple generation, relying either on excitation or inhibition as the main pacemaker. Excitation-first models assume that sharp wave and ripple oscillation are generated jointly by the sparsely recurrent pyramidal cell network, either via axonal gap junctions and antidromic spike propagation (Traub et al., 1999; Traub and Bibbig, 2000) or via supralinear dendritic integration (Memmesheimer, 2010; Jahnke et al., 2015). More recently, a GABA_A_ receptor-mediated excitatory feedback loop between axo-axonic chandelier interneurons and pyramidal cells has been proposed as a generating mechanism for SPW-Rs in the basolateral amygdala (Perumal et al., 2021).

Inhibition-first models focus on the generation of the ripple and posit that the interneuron network (e.g. in CA1) produces this fast oscillation in response to transient excitatory input due to a sharp wave event, which is generated by a separate mechanism upstream (e.g. in CA3) (Evangelista et al., 2020; Levenstein et al., 2019; Ecker et al., 2022). Inhibition-first models can be further subdivided into two classes, which we will call here the perturbation-based (Malerba et al., 2016) and the bifurcation-based inhibitory model (Buzsáki et al., 1992; Ylinen et al., 1995; Brunel and Hakim, 1999; Brunel and Wang, 2003; Donoso et al., 2018). The perturbation-based model assumes that the sharp wave-associated excitatory drive triggers a transient ringing effect in the interneuron population activity: While the activity settles into a new steady state, it produces a transient ripple of a characteristic duration that is determined by the amount of noise and heterogeneity in the network (Malerba et al., 2016). The bifurcation-based model, on the other hand, assumes that the interneuron network undergoes a bifurcation into a state of self-sustained oscillations (Brunel and Hakim, 1999), thus the ripple is only terminated due to the transient nature of the sharp wave-associated drive.

Experiments have remained inconclusive as to which of the proposed mechanisms is the most plausible for ripple generation (Maier and Kempter, 2017; Buzsáki, 2015). Previous analyses have focused on the average frequency and duration of ripple events as well as their dependency on pharmacological or optogenetic manipulation (Schlingloff et al., 2014; Stark et al., 2014; Buzsáki, 2015; Donoso et al., 2018). We propose that taking into account the *transient* dynamical features of spontaneous ripples can advance model selection. The instantaneous ripple frequency has been shown to decay over the course of a ripple event. This intra-ripple frequency accommodation (IFA) has been observed both *in vivo* and *in vitro* and across different animals (rat, mouse), brain states (sleep, awake), and measurements (LFP, inhibitory postsynaptic currents) (Ponomarenko et al., 2004; Nguyen et al., 2009; Sullivan et al., 2011; Hulse et al., 2016; Stark et al., 2014; Donoso et al., 2018).

So far, only the bifurcation-based inhibitory ripple model has been shown to reproduce IFA (Donoso et al., 2018). Here we use a theoretical mean-field approach and numerical simulations to explain the mechanism of IFA, show that it is robust with respect to parameter variation, and predict that it depends solely on the slope of the external, sharp wave-associated drive.

## Results

The bifurcation-based inhibitory ripple model assumes that a network of synaptically coupled CA1 interneurons, such as PV^+^ basket cells, acts as a delayed negative feedback loop and thus creates fast ripple oscillations when stimulated with excitatory, sharp wave-associated drive from CA3 (Brunel and Hakim, 1999; Donoso et al., 2018, see also Buzsáki, 1986; Ylinen et al., 1995; Brunel and Wang, 2003; Taxidis et al., 2012). Simulations of a biophysically detailed version of this model by Donoso et al. (2018) revealed that it can reproduce intra-ripple frequency accommodation (IFA) in response to time-dependent, sharp wave-like drive.

To explain the mechanism of IFA analytically, we first demonstrate that IFA is preserved in a spiking network model of reduced complexity (cf. Brunel and Hakim, 1999). We then turn to a mean-field approximation of the network dynamics, which enables us to study the ripple dynamics and IFA as a function of the time course of the excitatory sharp wave-like drive.

### Ripples and IFA in a spiking neural network model

We model the CA1 interneuron network as a homogeneous network of *N* leaky integrate-and-fire (LIF) neurons with time constant *τ_m_*, capacitance *C*, and resting potential *E*_leak_. The membrane potential *v_i_* of a unit *i* is given by the following stochastic differential equation:

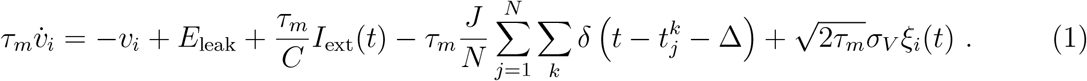

Whenever the membrane potential crosses a spike threshold *V*_thr_, a spike is emitted and the membrane potential is reset instantaneously to a reset potential *V*_reset_. For simplicity there is no absolute refractory period. All interneurons receive the same external, excitatory drive *I*_ext_ and an independent Gaussian white noise input *ξ_i_*, with zero mean ⟨*ξ_i_*(*t*)⟩ = 0 and unit noise intensity ⟨*ξ_i_*(*t*)*ξ_j_*(*t*′)⟩ = *δ_ij_δ*(*t* − *t*′), scaled by the noise strength parameter *σ_V_*. The network is fully connected via inhibitory pulse coupling of strength *J* and with a synaptic delay Δ (see Methods for details and default values of parameters).

The empirical population activity *r_N_*(*t*) in a small time interval [*t, t* + Δ*t*) is defined as the number of spikes *n*_spk_(*t, t* + Δ*t*) emitted by the population (cf. Methods, Eq. (15)), divided by the size of the population and the time step Δ*t*:

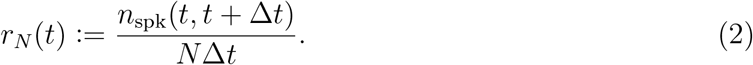

For plotting purposes *r_N_* is smoothed with a narrow Gaussian kernel (Methods, Eq. (16)). In the following we will illustrate that the key network dynamics in response to constant drive *I*_ext_ and time-dependent, sharp wave-like drive *I*_ext_(*t*), as presented by Donoso et al. (2018), are preserved in this model, *i.e*., there are fast oscillations in the ripple range and there is IFA.

The dynamics for constant drive are illustrated in Fig. 1. At low drive *I*_ext_ the network is in a steady-state with units firing asynchronously and irregularly at an overall low rate *f*_unit_ (Fig. 1A, left). As the drive increases, the network activity begins to exhibit coherent oscillations (Fig. 1A, middle). In the following, we will refer to the dominant frequency of this population oscillation as the network frequency *f*_net_ (Methods, black triangle markers in Fig. 1B). For a biologically reasonable set of parameters (Methods) the network frequency lies within the ripple range for a large range of external drives (Fig. 1B, gray band). The unit activity underlying the network ripple oscillation is sparse and irregular, *i.e*. the average unit firing rate *f*_unit_ is lower than the frequency *f*_net_ of the network oscillation (*saturation s* = *f*_unit_/*f*_net_ < 1) and the coefficient of variation (CV) of interspike intervals is around 0.5 (Fig. 1B, bottom panels). We hence refer to this state as *sparse synchrony* (Brunel and Hakim, 2008; Donoso et al., 2018).

**Figure 1:**
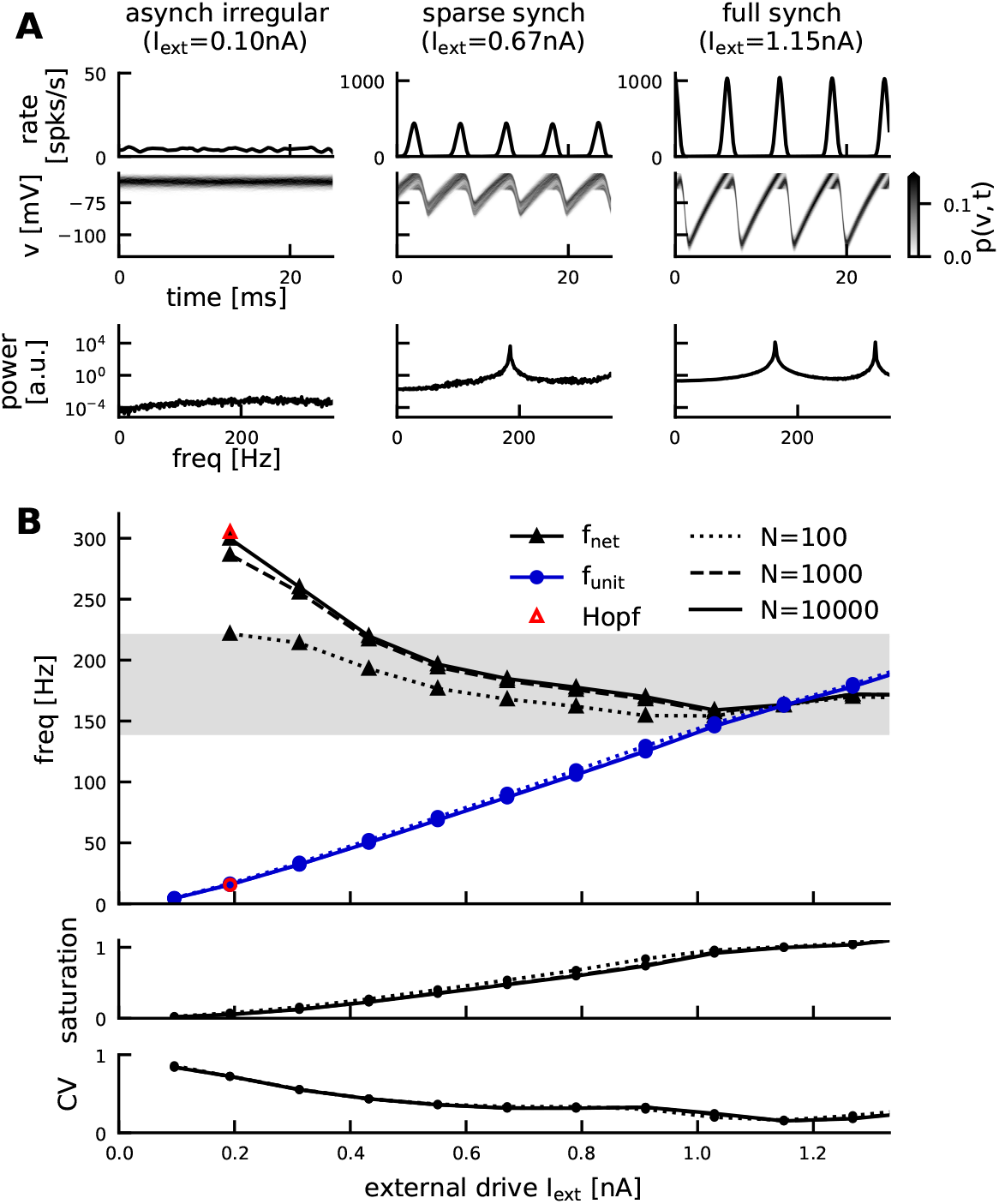
Constant-drive dynamics of the spiking network. **A**, Three dynamical regimes depending on the external drive: asynchronous irregular state, sparse synchrony, full synchrony (left to right, *N* = 10, 000). Top: population rate *r_N_*, middle: histogram of membrane potentials *v* (normalized as density), bottom: power spectral density of the population rate. **B**, Top: network frequency *f*_net_ (black) and unit firing rate *f*_unit_ (blue, average 1 SD) for a range of constant external drives *I*_ext_. Grey band marks approximate ripple frequency range (140– 220 Hz). Red markers indicate the critical input level 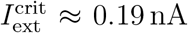 (Hopf bifurcation) and the associated network and unit frequency, as resulting from linear stability analysis (Brunel and Hakim, 1999, see Methods, Eq. (25)). Linestyle indicates network size (*N* ∈ [10^2^, 10^3^, 10^4^]). Middle: saturation *s* = *f*_unit_/*f*_net_. Bottom: coefficient of variation of interspike intervals.

In the mean-field limit *N* → ∞, the transition to this oscillatory state is sharp (a supercritical Hopf bifurcation) and can be determined by a linear stability analysis of the stationary state (Brunel and Hakim, 1999). In a simulated spiking network of finite size, the transition to the oscillatory state is not perfectly sharp because of inevitable fluctuations. Nevertheless, at the bifurcation point, both the network frequency and the unit firing rate are well predicted by the mean-field theory if the network is large enough (Fig. 1B, red markers).

For some sufficiently strong drive, the network reaches a state of full synchrony (*s* ≈ 1), with units firing regularly and at the same average frequency as the population rhythm (Fig. 1A, right). If the drive increases beyond this level, the network *de*synchronizes, since units start spiking several times per cycle (*s* > 1). Without an absolute refractory period, this desynchronization effect happens immediately after full synchrony is reached (cf. Donoso et al., 2018). Since the firing rates in that regime are too high to be biologically plausible, we focus in the following on the dynamical regime between the onset of oscillations (bifurcation point) and the point of full synchrony (Fig. 1).

During a sharp wave (SPW) event, which is thought here to be generated in CA3 (cf. models introduced by Evangelista et al., 2020; Levenstein et al., 2019; Ecker et al., 2022), pyramidal cells in CA3 transiently increase their firing rates (Csicsvari et al., 1999; Stark et al., 2014). Here we model the resulting feedforward input *I*_ext_(*t*) to the CA1 interneuron network in its simplest form: as a symmetric, piecewise linear double-ramp (Fig. 2A, bottom; see also Methods, Eq. (17)). This SPW-like drive elicits a transient ripple event in the network (Fig. 2A). We measure the instantaneous frequency of the population activity, either by taking a windowed Fourier transform (wavelet spectrogram) and finding the first peak in power in each time step, or by taking the inverse of the distances between consecutive peaks in the population activity providing frequencies for a discrete set of time points (Fig. 2A, top, continuous vs. discrete estimate). With both estimation methods, the instantaneous frequency shows an asymmetry: frequencies in the first half of the event are clearly higher than in the second half (Fig. 2A, top, see also Donoso et al., 2018), *i.e*. the network exhibits intra-ripple frequency accommodation (IFA).

**Figure 2:**
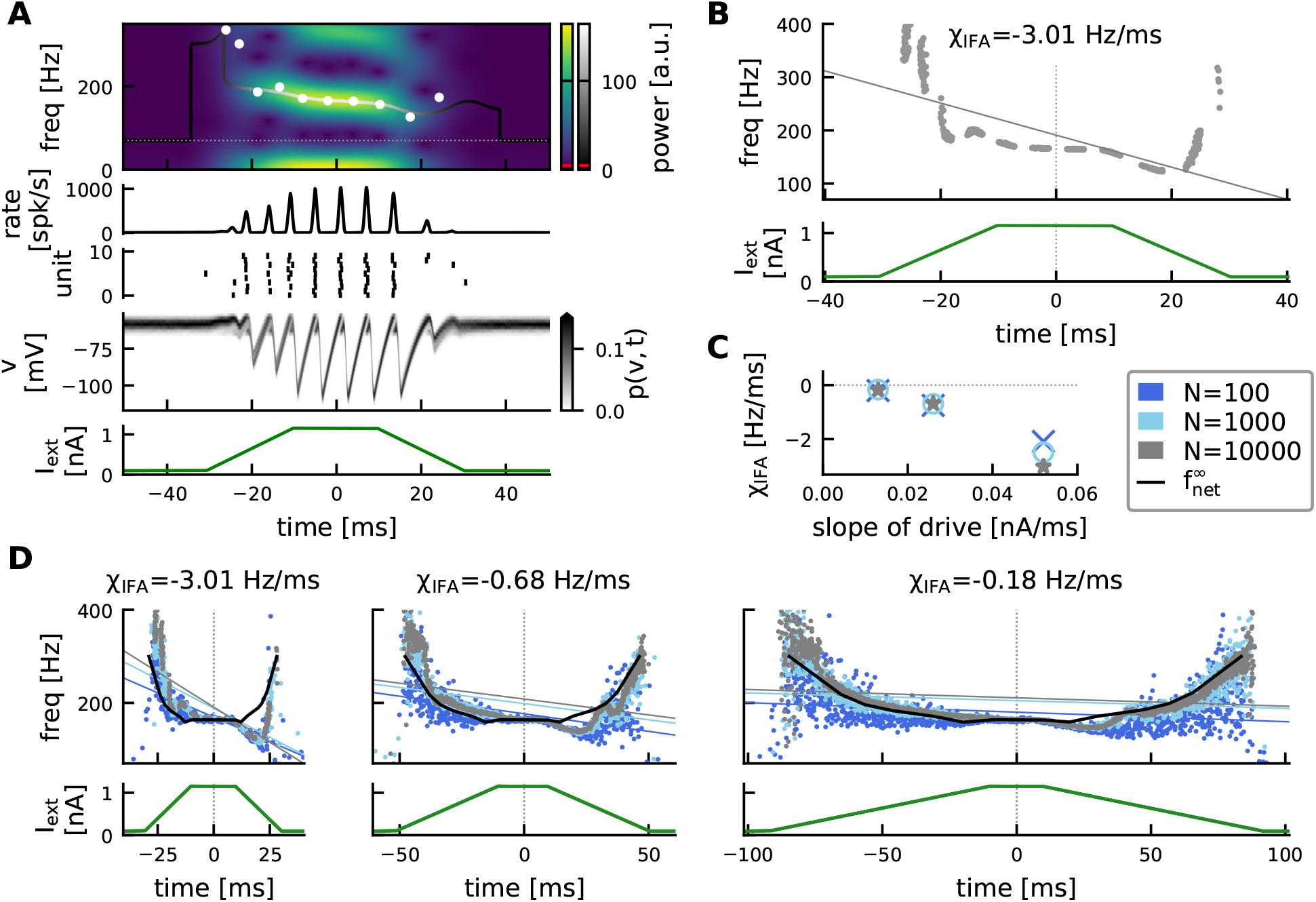
Transient dynamics of the spiking network and IFA. **A**, Example simulation showing a transient ripple with IFA. From bottom to top: SPW-like drive (Eq. (17)); histogram of membrane potentials (normalized); raster plot showing spike times of 10 example units; population rate exhibiting transient ripple oscillation; wavelet spectrogram indicating instantaneous power (blue-yellow colorbar) for a frequency range of 0–350 Hz. Solid curve: continuous estimate of instantaneous frequency based on wavelet spectrogram with gray scale indicating maximal instantaneous power. Dotted line: cutoff frequency for peak detection *f*_min_ = 70 Hz. Red line: power threshold (see Methods). White dots: discrete estimate of instantaneous frequency based on peak-to-peak distance in population rate. Network size *N* = 10, 000. **B**, Quantification of IFA. Top: Grey dots: discrete instantaneous frequency estimates from 50 independent simulations. Grey line: linear regression line with negative slope (*χ*_IFA_ = −3.01 Hz/ ms) indicating IFA (see Methods, Eq. (18)). Bottom: The same SPW-like drive was applied in all 50 simulations. Network size *N* = 10, 000. **C**, Dependency of IFA slope *χ*_IFA_ on the slope of the external drive for different network sizes (color coded). **D**, Instantaneous (dots) vs asymptotic (black line) network frequencies (top) for piecewise linear drives (bottom) of decreasing slopes (left to right). Color indicates network size *N*. Thin, colored linear regression lines illustrate decreasing strength of IFA for shallower drive. The indicated IFA slope *χ*_IFA_ refers to the largest network (*N* = 10, 000). With decreasing slope of the drive (from left to right) the instantaneous network frequencies become more similar to the symmetric, asymptotic frequencies and IFA vanishes. Network size does not influence the IFA slope significantly (see also C). Asymptotic network frequencies are derived via interpolation of the constant-drive results shown in Fig. 1.

There is no rigorous definition of IFA. The central aspect of IFA is the decrease (*accommodation*) of the instantaneous frequency over the course of a ripple event. We observe such a decrease in the central portion of this simulated ripple event, where power in the ripple band is large (Fig. 2A, top). In the model, this decay is not strictly monotonic over the entire event. We observe a small increase of the frequency at the end of the event, albeit with low power (the underlying peaks in the population rate are small). Similar non-monotonic trajectories of the ripple frequency have been observed in experimentally measured ripples and still been classified as exhibiting IFA (Ponomarenko et al., 2004). Hence, here we define IFA not as a strictly monotonic decay, but as a more general trend from higher towards lower frequencies over the course of an event. Accordingly, we quantify the IFA asymmetry by linear regression over the discrete instantaneous frequency estimates. This quantification of IFA can also be applied to results from many simulations with different noise realizations and the same SPW-like drive (Fig. 2B; Methods, Eq. (18)). A negative regression slope (here *χ*_IFA_ = −3.01 Hz/ ms) indicates IFA.

In the model, the IFA asymmetry can be understood best by comparing the instantaneous frequencies to the asymptotic frequencies that the network would settle into if the drive remained constant at any given level indefinitely (Fig. 2D, black lines). Naturally, these asymptotic frequencies follow the same symmetry as the external drive and thus provide a useful reference frame. For our double-ramp drive (Methods, Eq. (17)), we observe that during the plateau phase (*i.e*. constant drive) the instantaneous frequency quickly approaches the asymptotic frequency. However, during the rising phase of the ramp-input the instantaneous frequency tends to be higher than the asymptotic reference. During the falling phase it is lower, thus creating the overall IFA asymmetry.

Varying the slope of the external double-ramp drive we see that the IFA asymmetry is speed-dependent (Fig. 2C,D): If the external drive changes more slowly (smaller slope), the network frequency response becomes more symmetric; for very small slopes the instantaneous frequencies approach the symmetric, asymptotic reference frequencies.

In what follows, we aim at understanding the mechanism behind the observations illustrated for constant drive in Fig. 1 and time-dependent drive in Fig. 2. Further simulations indicate that the network dynamics varies only little when the network size is varied by two orders of magnitude (Fig. 1B, Fig. 2C,D). We hence hypothesize that IFA is preserved in the mean-field limit of an infinitely large network, and will use a mean-field approach to explain the generating mechanism of IFA.

### Gaussian-drift approximation of ripple dynamics in the mean-field limit

To facilitate notation in the mathematical analysis, we rescale all voltages to units of the distance between threshold and rest such that the new spiking threshold is at *V_T_* = 1 and the resting potential is at *E_L_* = 0 (cf. Methods, Eq. (19)). The single unit stochastic differential equation (previously Eq. (1)) then reads

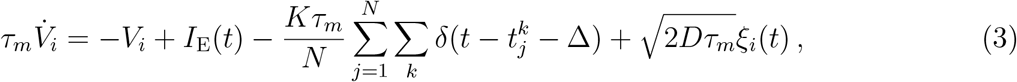

with rescaled external excitatory current *I*_E_, inhibitory synaptic strength *K*, and noise intensity *D* (Methods, Eq. (21)). In the mean-field limit *N* → ∞, the dynamics of the density of membrane potentials *p*(*V, t*) is described by the Fokker-Planck equation (FPE)

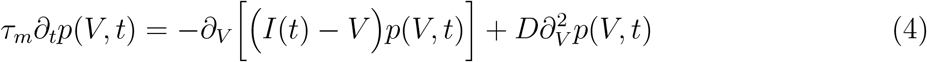

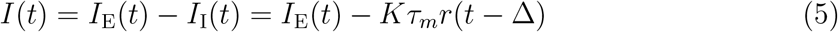

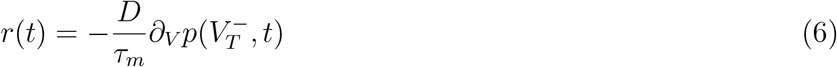

(see Methods, Eqs. (22) for details). The population rate *r*(*t*) defined in Eq. (6) represents the mean-field limit of the population activity *r_N_*(*t*) in Eq. (2) of the spiking neural network. Compared to the classical application of the FPE (Risken, 1989; Gardiner, 1985) there are two essential differences: First, the FPE is nonlocal because of the resetting rule. This rule imposes an absorbing boundary condition at the threshold (Eq. (22e)) and a source of probability at the reset point; the latter source can be imposed by a jump condition for the derivative of the density at the reset point (Eq. (22g)) that matches the derivative at the threshold point (see *e.g*. Abbott and van Vreeswijk, 1993; Brunel, 2000; Lindner and Schimansky-Geier, 2001; Fourcaud and Brunel, 2002; Brunel et al., 2003). Secondly, the FPE (4) is nonlinear because the current *I*(*t*) depends on the probability density *p*(*V, t*) through the population rate *r*(*t*) in Eq. (6).

Stable stationary solutions of this nonlocal, nonlinear FPE correspond to asynchronous irregular spiking (Abbott and van Vreeswijk, 1993; Brunel and Hakim, 1999). It has been shown that an oscillatory (*i.e*. periodic) solution *r*(*t*) emerges via a supercritical Hopf bifurcation when the external drive *I*_E_ exceeds a critical value (Abbott and van Vreeswijk, 1993; Brunel and Hakim, 1999). This network oscillation well reproduces the coherent stochastic oscillation of the population activity *r_N_* (*t*) in the finite-size spiking neural network (Fig. 1). The network frequency at the onset of oscillations (*i.e*. at the Hopf bifurcation, where the stationary solution looses stability) is well predicted analytically by a linear stability analysis (Brunel and Hakim, 1999, see also Methods, Eq. (25) and Fig. 1B, red markers). However, this analytical prediction quickly breaks down further away from the bifurcation where the oscillation dynamics becomes strongly nonlinear (see power spectral densities with higher harmonics in Fig. 1A). In this regime neither an exact periodic solution of the FPE nor an approximation of its frequency is known. We will thus introduce a simplified approach that allows us to approximate the network frequencies at strong drive beyond the bifurcation.

Our approach can be motivated by two observations from the spiking network simulations (Fig. 1): (a) In the relevant regime between sparse and full synchrony, units spike at most *once* per cycle of the population rhythm (*f*_unit_ ≤ *f*_net_, Fig. 1B). This property will allow us to approximate the time course of a population spike in *r_N_* (*t*) using a first-passage-time ansatz, neglecting the reset mechanism. (b) In between population spikes, the population rate *r_N_* (*t*) is close to zero, and the strong inhibitory feedback pushes the bulk of the membrane potential distribution significantly below threshold (Fig. 1A). In those periods the absorbing boundary condition at threshold does not have any significant impact on the dynamics of *p*(*V, t*).

These two observations motivate a considerable simplification: Without the boundary conditions at threshold and reset, the FPE (4) can be solved analytically. In the long-time limit its solution becomes a simple Gaussian — independent of the initial condition (Methods, Eq. (26)). We thus approximate the density of membrane potentials as

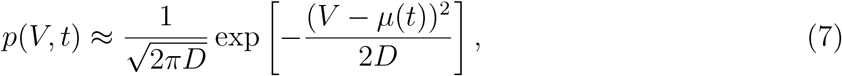

where the only time-dependent quantity is the mean membrane potential *μ*(*t*), which evolves according to

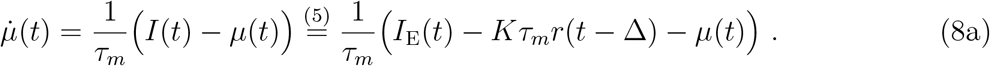

Because in the considered regime spikes are mainly driven by the mean input rather than membrane potential fluctuations, the population rate can be well approximated by the drift part of the probability current across the threshold, while diffusion-mediated spiking is ignored (Methods, Eqs. (30) and (31); see also Goedeke and Diesmann, 2008; Plesser and Gerstner, 2000; Chizhov and Graham, 2007; Schwalger, 2021):

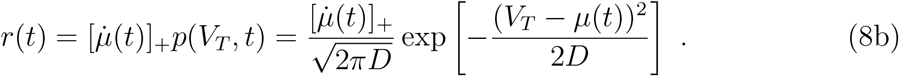

Thus, in our approximate dynamics without reset, the population rate *r*(*t*) is given by the membrane potential density at threshold, scaled by the speed 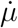 at which the mean membrane potential approaches the threshold. Whenever the mean membrane potential is *de*creasing 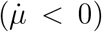, the drift current at the threshold is negative, and hence, the rate is clipped to 0 ([*x*]_+_ ≔ (*x* + *x*)/2).

The system of two coupled equations, Eqs. (8), is equivalent to a single delay differential equation (DDE; Methods, Eq. (33)). This DDE governs the dynamics of the mean of the Gaussian membrane potential distribution, and the resulting drift-based population rate. In the following, we will therefore refer to the DDE dynamics as the *Gaussian-drift approximation*.

The main differences between the spiking network model (or the exact mean-field dynamics given by the FPE (4)) and the Gaussian-drift approximation are illustrated in Fig. 3 for the case of constant drive. In the spiking network model, the membrane potential density *p*(*V, t*) is not strictly Gaussian. Because of the fire-and-reset mechanism, *p*(*V, t*) changes in shape during the oscillation cycle, becoming at times even bimodal (Fig. 3Aiii vs Biii). Still, we see that our simplifying assumption (b) is justified: Whenever the membrane potential density is subthreshold in between population spikes, it becomes more Gaussian, and its standard deviation approaches 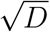 (Fig. 3 Aii, bottom; Aiii, first 3 snapshots).

**Figure 3:**
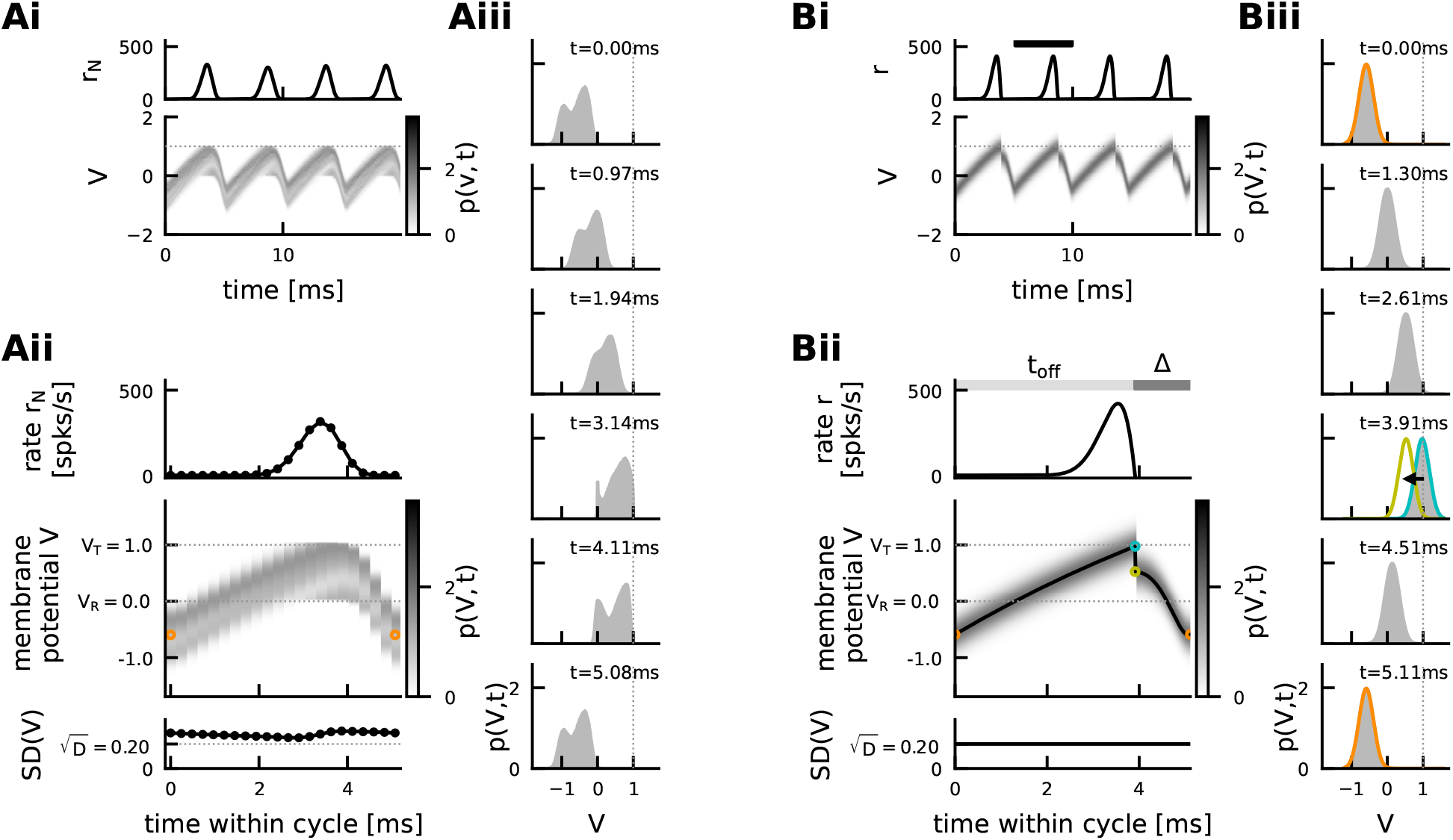
Illustration of the Gaussian-drift approximation. Comparison of oscillation dynamics in the spiking network simulation (A) and the Gaussian-drift approximation (B) at constant drive (*I*_E_ = 4.24). **Ai**, Spiking network simulation. Empirical population rate *r_N_* (top), and density of membrane potentials (bottom), exhibiting coherent stochastic oscillations with (weak) finite size fluctuations. Dotted line marks spike threshold *V_T_* = 1. **Aii**, The average cycle of the oscillation dynamics in Ai (computed for 21 bins of 0.24 ms each). Top: population rate; middle: density of membrane potentials, orange marker: local minimum of mean membrane potential; bottom: standard deviation of membrane potential distribution, dotted line: theoretical asymptote 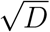 in the absence of boundary conditions (Eq. (28)). **Aiii**, Snapshots of the membrane potential density over the course of the average cycle shown in (Aii). Dotted line marks spike threshold/absorbing boundary. **Bi**, Gaussian-drift approximation (Eqs. (8)). Top: population rate, bottom: Gaussian density of membrane potentials *p*(*V, t*), both perfectly periodic. **Bii**, Zoom into one oscillation cycle (black bar in Bi). Top: population rate *r*, middle: density of membrane potentials *p*(*V, t*), bottom: constant standard deviation 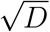. The mean membrane potential *μ*(*t*) (black line) starts each cycle in *μ*_min_ (orange) and rises up until *μ*_max_ (cyan) at time *t*_off_, at which point the population spike ends. In a phenomenological account for the single unit reset, *μ* is reset instantaneously to *μ*_reset_ (yellow, Eq. (51)). From there *μ* declines back towards *μ*_min_. **Biii**, Snapshots of the membrane potential density *p*(*V, t*) over the course of one cycle (Bii). Colors mark *t* = 0 ∼ *T* (orange), and *t* = *t*_off_ (cyan/yellow). Note that in the theoretical approximation the spiking threshold *V_T_* = 1 (dotted line) is no longer an absorbing boundary.

The advantage of our Gaussian-drift approximation is that it reduces the FPE with complex boundary conditions to a simpler DDE with oscillatory solutions that can be studied analytically. In the following, we will first consider the dynamics for constant drive, then extend the Gaussian-drift approximation to understand the transient response to time-dependent drive, and hence explain the emergence of IFA.

### Analysis of oscillation dynamics for constant drive

A *T*-periodic solution of the Gaussian-drift model, Eqs. (8), must have a mean membrane potential *μ*(*t*) = *μ*(*t* + *T*) that oscillates between two local extrema *μ*_min_ and *μ*_max_ (Fig. 3Bi). Whenever the mean membrane potential *in*creases, a positive population rate is produced; when the mean membrane potential *de*creases the population rate is clipped to 0 (Eq. (8b)). Let us consider a single cycle of this oscillatory dynamics (Fig. 3Bii): The moment when *μ* reaches its local maximum *μ*_max_ is of special importance as it marks the end of the population spike. We will refer to this time as *t*_off_:

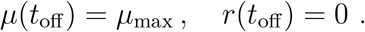

Since the inhibitory feedback *I*_I_(*t*) = −*Kτ_m_r*(*t* − Δ) is proportional to the population rate with a delay of Δ (Eq. (8a)), it follows that this feedback ceases exactly Δ after the end of the population spike, *i.e. I*_I_(*t*_off_ + Δ) = 0. It is convenient to define this time as the end of a cycle, *i.e*. the beginning of the next one. The mean membrane potential at this time is close to its local minimum (see Methods, Step 2) and will be denoted as *μ*_min_ ≔ *μ*(*t*_off_ + Δ). The period *T* can then be split into the *upstroke* time *t*_off_ needed for the mean membrane potential to rise from *μ*(*t* = 0) = *μ*_min_ towards *μ*(*t*_off_) = *μ*_max_, and the *downstroke* time Δ during which the mean membrane potential is pushed back down to *μ*(*T*) = *μ*_min_ due to the delayed inhibitory feedback (Fig. 3Bii). In Methods we derive, through a series of heuristic approximations, analytical expressions for the local extrema *μ*_max_ (Eq. (40)) and *μ*_min_ (Eq. (52)) of the mean membrane potential oscillation as a function of the external drive *I_E_*. Using these expressions, we obtain an analytical formula for *t*_off_ (Eq. (49)) and hence for the network frequency:

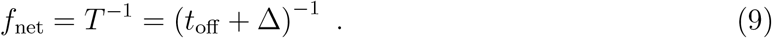

Apart from the network frequency the Gaussian-drift approximation also allows an intuitive understanding of the mean unit firing rate. When the population spike ends at time *t*_off_, the suprathreshold portion of the Gaussian density corresponds to the fraction of units that have spiked in the given cycle (the *saturation s*, Methods, Eq. (50)). The mean unit firing rate can thus be inferred as:

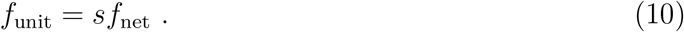

In the following we compare our analytical Gaussian-drift approximations for the network frequency and mean membrane potential dynamics to the spiking network simulations for a range of external drives *I*_E_.

### Evaluation of the performance of the Gaussian-drift approximation

Our theory captures the dependence of the network frequency *f*_net_ and the mean unit firing rate *f*_unit_ on the external drive *I*_E_, including the transition from sparse to full synchrony for increasing external drive (Fig. 4, top). Our analytically derived expression for the saturation *s* predicts this transition (Fig. 4, middle), since *s* is a monotonically increasing function of *μ*_max_ (Eq. (50)), which monotonically increases as a function of the external drive *I*_E_ (Eq. (40), Fig. 4, bottom). We can also estimate analytically the point of full synchrony, *i.e*. the amount of external drive 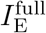 that is required for single units to fire approximately at the frequency of the network rhythm 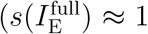; Methods, Eq. (53); Fig. 4, vertical dashed line). The Gaussian-drift approximation slightly overestimates the point of full synchrony, but correctly predicts its parameter dependencies: For stronger coupling and/or larger noise, stronger external drive is required to reach full synchrony (Supplementary Fig. S2).

**Figure 4:**
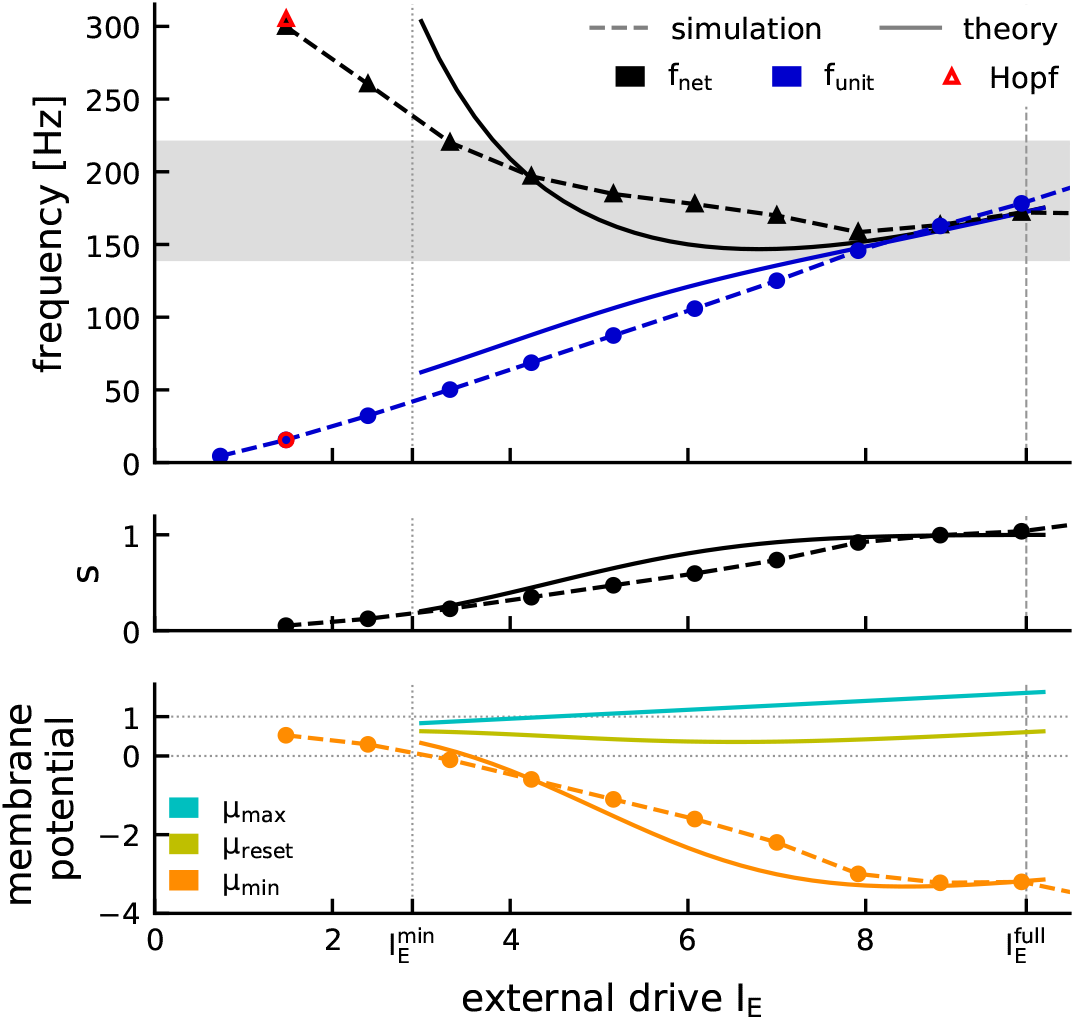
Analytical approximation of the oscillation dynamics for constant drive. Comparison of dynamics in theory (full lines) and spiking network simulation (dashed lines). Top: Network frequency (black triangles) and mean unit frequency (blue circles). Red markers: Hopf bifurcation. Vertical lines indicate the range 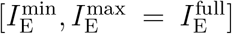 for which the theory applies (see Methods, Eq. (54)). Middle: Saturation *s* increases monotonically with the drive (Eq. (50)). Bottom: characterization of the underlying mean membrane potential dynamics via local maximum *μ*_max_ (cyan, Eq. (40)), local minimum *μ*_min_ (orange, Eq. (52)) and population reset *μ*_reset_ (yellow, Eq. (51)). Default parameters (see Methods).

The theory shows that the amplitude *μ*_max_–*μ*_min_ of the oscillatory mean membrane potential grows with increasing drive (Fig. 4, bottom). This is mainly due to a strong decrease of the periodic minimum *μ*_min_ (Fig. 4, bottom, solid orange line), which we also observe in the spiking network simulation (Fig. 4, bottom, dashed orange line). The quantities *μ*_max_ and *μ*_reset_ are pertinent to the Gaussian-drift approximation and have no direct counterpart in the spiking neural network model.

The range of applicability of our theory is defined by our two assumptions (see Methods for details, Eq. (54)): (a) units should spike at most once per cycle (*i.e.* 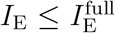), (b) in between population spikes the bulk of the membrane potential distribution should be subthreshold (*i.e.* 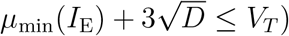). The resulting range 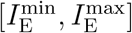 for the external current *I*_E_ covers the large part of the regime of sparse synchrony up to the point of full synchrony (Fig. 4). We confirmed with numerical simulations that for strong enough drive the Gaussian-drift approximation works for a wide parameter regime w.r.t. noise, coupling strength, and synaptic delay (see Supplement, Fig. S1, Fig. S2).

At low drive 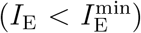 this theory breaks down (see also Methods, *Numerical analysis of oscillation dynamics*). This is to be expected from a purely drift-based approximation. The dynamics of the spiking network close to its supercritical Hopf bifurcation is largely fluctuation-driven. Such dynamics cannot be captured by focusing only on the oscillation of the mean membrane potential, which has infinitesimal amplitude as the drive approaches its critical value. This limitation of the theory does not pose a problem, since (a) the fluctuation-driven dynamics around the Hopf bifurcation has already been studied in depth by Brunel and Hakim (1999) and (b) our main goal here is to explain the IFA dynamics, which happens in the strongly mean-driven regime 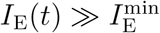.

So far we have studied the case of constant drive describing the *asymptotic* oscillatory dynamics observed in the long-time limit, after initial conditions have been forgotten. This will be emphasized from here on by adding a superscript “∞”. We will now study how the *transient* dynamics, introduced either by a perturbation of the initial condition or a time-dependent drive, deviate from these asymptotic constant-drive dynamics. We will demonstrate that the strong monotonic increase of the asymptotic oscillation amplitude with increasing values of the constant drive can lead to a speed-dependent hysteresis effect and IFA under time-dependent drive.

### Analysis of oscillation dynamics for piecewise constant, sharp wave-like drive

Even for constant drive, there is *transient* dynamics if the initial mean membrane potential deviates from the asymptotic minimum 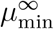. Let us assume that a cycle starts with an initial mean membrane potential 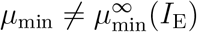. We only require that *μ*_min_ is sufficiently subthresh-old, such that the initial population rate is close to zero. What will be the period of the first cycle and how long does it take until the asymptotic dynamics is reached?

First we note that, independent of *μ*_min_, the mean membrane potential will rise towards the asymptotic 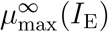, which can be shown to be independent of the initial condition (Methods, Eq. (40)). Thus, only the duration of the first upstroke will be influenced by the initial condition, and the asymptotic dynamics is reached immediately thereafter.

The duration of the first upstroke depends on the distance that the mean membrane potential has to travel, from its initial value *μ*_min_ to the next peak 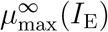. For 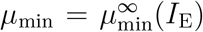 the upstroke has length 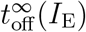 and hence the length of the first cycle is equal to the asymptotic period *T*^∞^. Correspondingly, the instantaneous frequency is equal to the asymptotic network frequency, 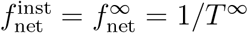 (Fig. 5A, middle). The upstroke takes *less* time, if the net net mean membrane potential starts at a *higher* value 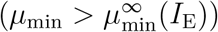 and more time otherwise (Fig. 5A, left vs right). Hence the period of the first cycle will be either shorter or longer, which can be rephrased as an instantaneous frequency 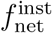 that is higher or lower than the asymptotic 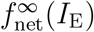. Fig. 5B illustrates the instantaneous frequency of the first cycle for different combinations of (constant) drive *I*_E_ and initial condition *μ*_min_ (red: instantaneous frequency is higher than asymptotic frequency; blue: instantaneous frequency is lower).

**Figure 5:**
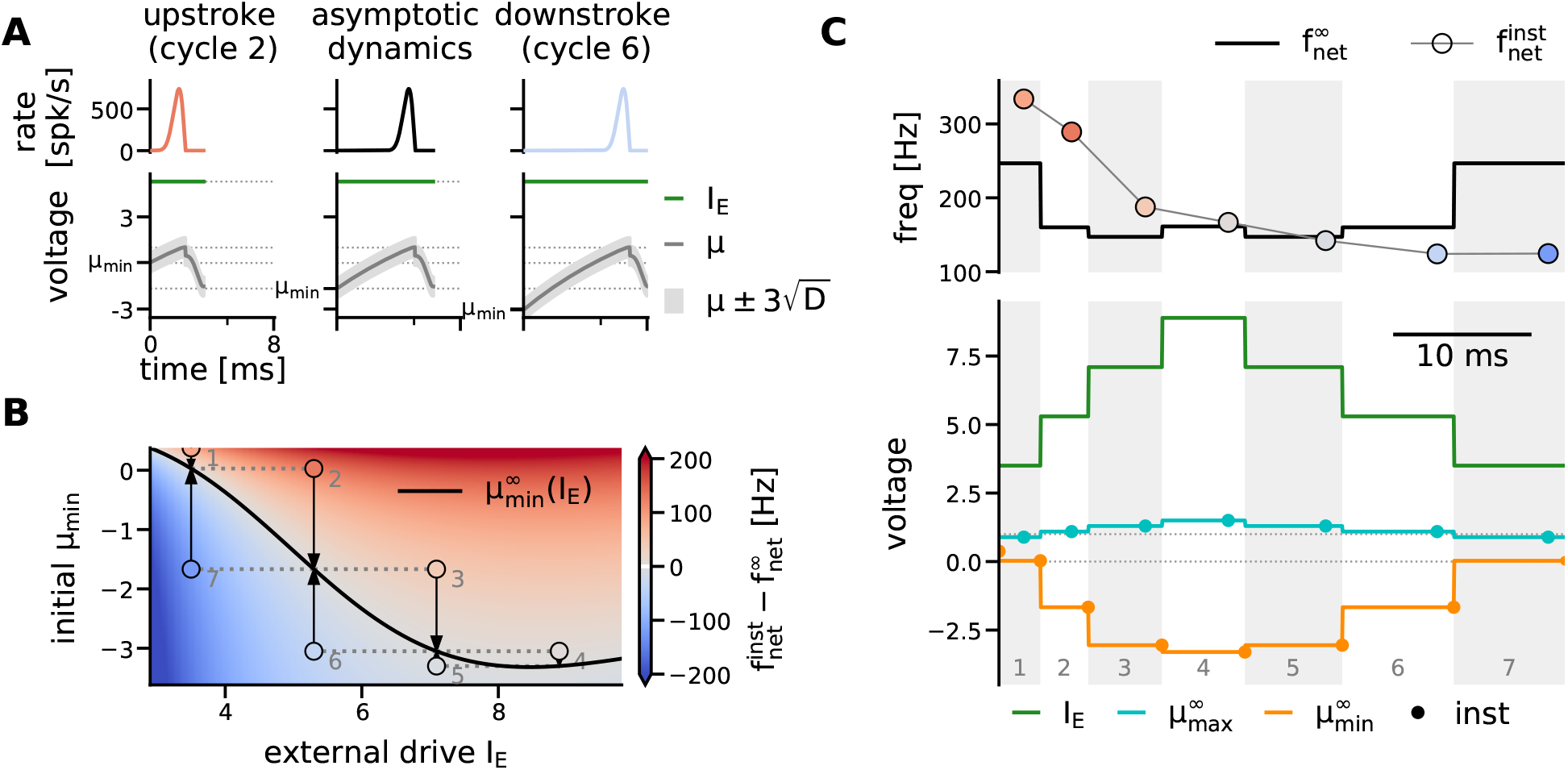
Transient dynamics and IFA for piecewise constant external drive. **A**, Dependence of the length of the oscillation cycle on the initial mean membrane potential. Left: shorter period for 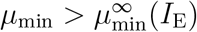 (cycle 2 in C). Middle: asymptotic period for 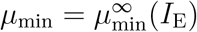. Right: longer period for 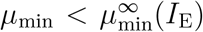 (cycle 6 in C). **B**, Difference between the instantaneous frequency of a cycle with constant drive *I*_E_ and initial condition *μ*_min_, and the asymptotic frequency 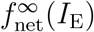 for a range of external drives *I*_E_ and initial mean membrane potentials *μ*_min_. Black line: asymptotic 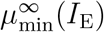 (cf. Fig. 4, bottom, orange line). Markers indicate example cycles shown in A, C. Arrows indicate convergence to the asymptotic dynamics after one cycle. If the drive changes after each cycle (dotted lines), the seven examples lead to the trajectory shown in C. **C**, IFA for piecewise constant drive with symmetric step-heights. Shaded areas mark oscillation cycles. Bottom: The external drive is increased step-wise, up to the point of full synchrony 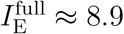 (green line). Lines in all panels indicate the asymptotic dynamics associated to the external drive of the respective cycle. Markers indicate transient behavior. Cyan: *μ*_max_ (reset not shown). Orange: *μ*_min_. Top: the instantaneous network frequency (markers) is first above and then below the resp. asymptotic network frequencies (black line).

Once the mean membrane potential has reached its first peak, it will follow the asymptotic dynamics, settling into 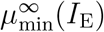 at the end of the first cycle, and all subsequent cycles will come at the asymptotic frequency 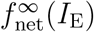 associated to the external drive *I*_E_ (convergence indicated by arrows in Fig. 5B). A change of initial condition can thus only introduce a transient deviation from the asymptotic dynamics in a single cycle.

What if we change the external drive after each cycle (green line in Fig. 5C)? Then the initial mean membrane potential of each cycle *i* will be the asymptotic minimum associated to the drive of the *previous* cycle:

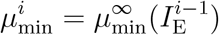

*i.e*. the mean membrane potential dynamics exhibits a history dependence (or *hysteresis*). Now recall that the asymptotic minimum 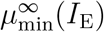 is a monotonically decaying function of the drive (except for strong drive close to 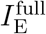, Fig. 5B, black line). Thus, if the external drive *in*creases stepwise, each cycle starts with an initial mean membrane potential *above* the asymptotic minimum associated to that cycle’s drive, hence the instantaneous frequency is *above* its asymptotic value (Fig. 5B, trajectory through red area: 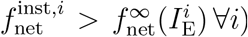. Vice versa, if the external drive *de*creases stepwise, each cycle starts with an initial mean membrane potential *below* the asymptotic minimum associated to that cycle’s drive, hence the instantaneous frequency is *below* its asymptotic value (Fig. 5B, trajectory through blue area: 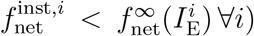. In summary:

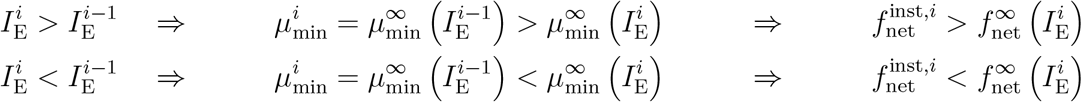

Thus, if we approximate the transient change in drive during a sharp wave as a simple, piecewise constant function that first increases after each cycle, and then decreases (Fig. 5C, green line), we observe IFA: The asymptotic network frequency associated to the drive in each cycle describes a reference curve that follows the same symmetry as the drive (Fig. 5C, top, solid black line). However, the instantaneous network frequency (Fig. 5C, top, round markers) is asymmetric over time, as it is *above* the asymptotic network frequencies during the rising phase of the external drive, and *below* during the falling phase. The theory thus describes the relationship between instantaneous and asymptotic frequencies that was already described for the spiking network simulations in Fig. 2D.

The piecewise constant shape of the drive may not be realistic, but serves to illustrate the core mechanism of IFA: a hysteresis in the oscillation amplitude of the mean membrane potential. A drawback is that this simple model for SPW-like drive is not symmetric in *time*, since the drive changes after each cycle, and the cycle length increases due to IFA. To show that the IFA asymmetry does not rely on an asymmetry in the drive, we adapted the Gaussian-drift approximation to incorporate time-dependent linear drive.

### Analysis of oscillation dynamics for piecewise linear, sharp wave-like drive

Following the same approach as before, we approximate the *transient* dynamics of the mean membrane potential in a cycle *i* with initial value 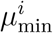 and a drive that changes linearly around a level 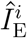 with slope *m*:

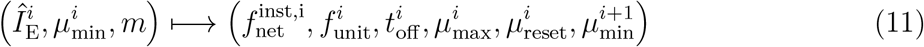

The dynamics is quantified in terms of the peak of the mean membrane potential 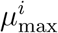, its reset value 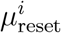, and the value 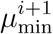 that is reached at the end of cycle *i* (and may thus be the initial membrane potential of the next cycle *i* + 1). Most importantly, the duration of the upstroke 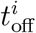 is inferred, and from that the instantaneous network frequency 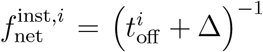 (see Methods, Eqs. (58)-(68)). In agreement with our theoretical approximation, which is anchored to the end of the population spike, the reference drive 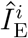 is chosen such that 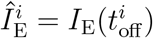, *i.e*.

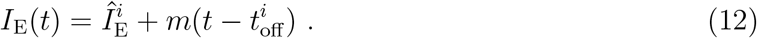

The analysis is now a little more complex, since each cycle depends on three parameters 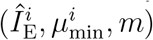, in contrast to only one parameter 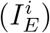 for piecewise constant drive. We will demonstrate, however, that the essential findings from the basic case of piecewise constant drive still hold, *i.e*. that IFA is generated by the same hysteresis in the transient dynamics of the mean membrane potential that we have uncovered before (Fig. 5 vs Fig. 6). To illustrate this by an example, we fix the slope of the linear drive to *m* = 0.4/ ms. We then compare how the transient dynamics deviates from the asymptotic dynamics, depending on whether the drive is increasing (*m* = +0.4/ ms) or decreasing (*m* = 0.4/ ms) (Fig. 6).

**Figure 6:**
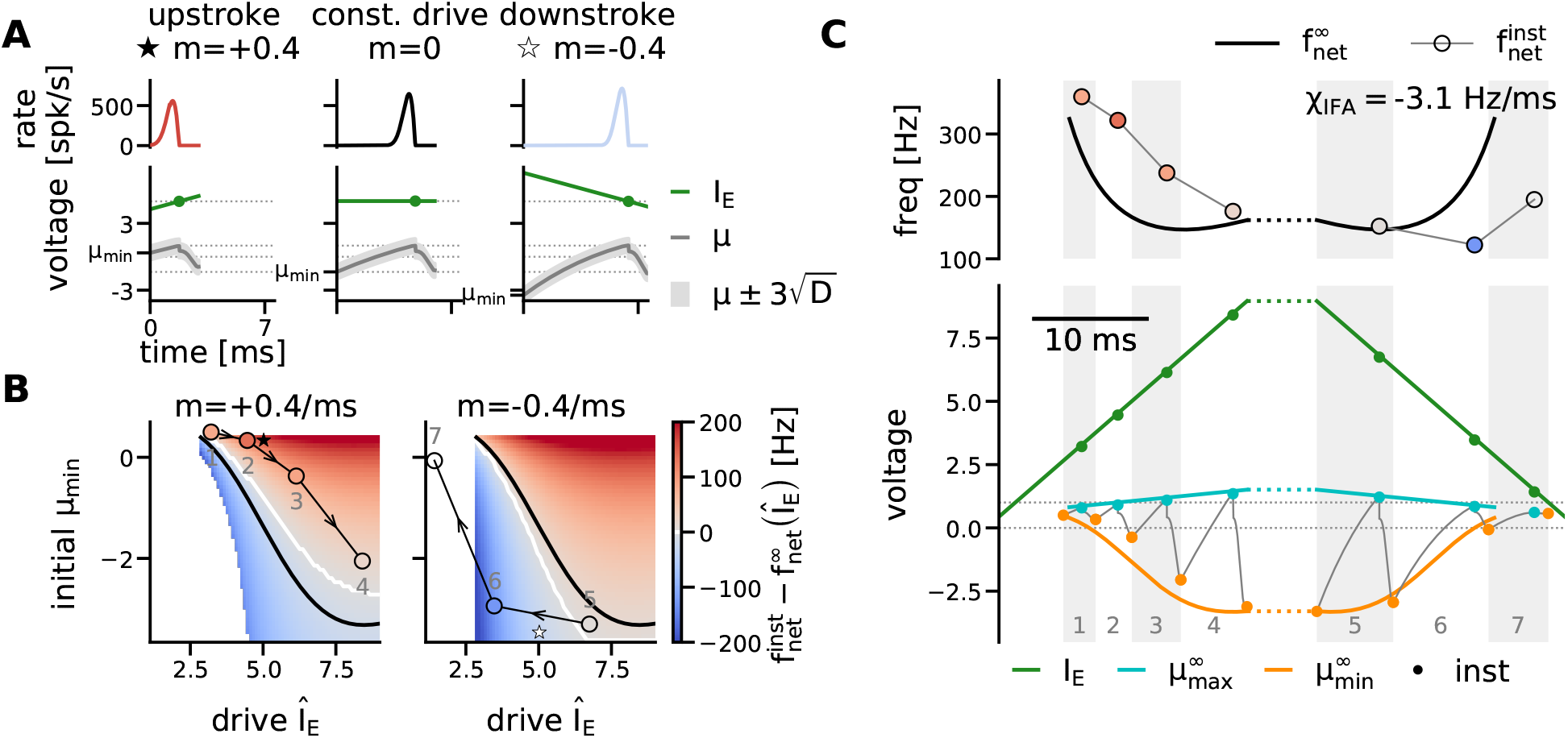
Transient dynamics and IFA for piecewise linear external drive. **A**, Exemplary transient dynamics during rising vs. falling phase of the external drive, given fixed *Î*_E_ = *I*_E_ (*t*_off_) = 5 (green dot). Left: shorter period for *m* > 0 and initial 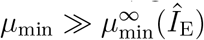. Middle: asymptotic period for constant drive (*I*_E_ ≡ *Î*_E_, *m* = 0) and initial 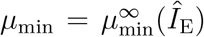. Right: longer period for *m* < 0 and initial 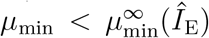. Dotted horizontal lines mark reference drive *Î*_E_, spike threshold *V_T_* = 1, reset potential *V_R_* = 0 and the asymptotic 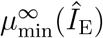. **B**, Difference between instantaneous and asymptotic frequency for a range of reference drives *Î*_E_ and initial mean membrane potentials *μ*_min_. Left: linearly increasing drive with slope *m* = +0.4/ ms. Right: linearly decreasing drive with slope *m* = −0.4/ ms. Black line: asymptotic 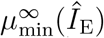 for constant drive. White line: initial membrane potential *μ*_min_, for which 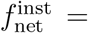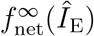. Stars mark the examples shown in A for *Î*_E_ = 5. Round markers and arrows indicate the trajectory shown in C for piecewise linear drive, numbered by cycle. White space where either: no asymptotic oscillations occur 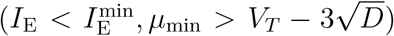, or (bottom left): no transient solution exists (see Methods, Eq. (66)). **C**, IFA for symmetric, piecewise linear (SPW-like) drive. Shaded areas mark oscillation cycles. Bottom: The external drive is increased up to the point of full synchrony 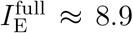 (green line). Colored lines indicate asymptotic dynamics. Thin grey line: mean membrane potential trajectory *μ*(*t*) in response to SPW-like drive *I*_E_(*t*) (numerical integration of DDE Eq. (8)). Markers quantify transient behavior. Cyan: *μ*_max_. Orange: *μ*_min_. Reset not shown to enhance readability. Top: the instantaneous network frequency (markers) is first above, then below the resp. asymptotic network frequencies (black line). Dashed lines: plateau phase of variable length with 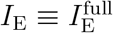, during which the network settles into the asymptotic dynamics.

The principal insight from the case of constant drive still holds for the case of time-dependent drive, except for a small range of initial values 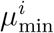: If the initial mean membrane potential 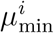 is (sufficiently) larger than the asymptotic reference 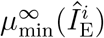, the period is shorter than the asymptotic reference *T*_∞_, hence 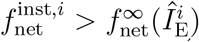 (Fig. 6A, left; B, red areas). In contrast, if 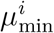 is smaller than the asymptotic reference 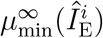, the period is longer than the asymptotic reference *T*^∞^, hence 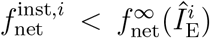 (Fig. 6A, right; B, blue areas). Exceptions from this “rule” (Fig. 6B, space between black and white line) occur because for time-dependent drive an asymptotic initial value 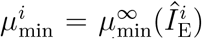 (black line) no longer implies the asymptotic period *T*^∞^ and frequency 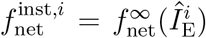 (white line, see Methods for details). We note, however, that these exceptions from the rule occur only in a small portion of the state space that is rarely visited in a given ripple event, as we will see in the following.

What can we say about the dynamics of consecutive cycles *i, i* + 1, … that occur if the drive rises or falls continuously with slope *m*? At the end of each cycle the mean membrane potential is close to the asymptotic reference 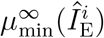 (Methods, Eq. (68)). Thus we observe the same hysteresis as before: if the drive *increases* (*m* > 0, Fig. 6B, left), trajectories of consecutive cycles will lie in the upper right half of the parameter space, where every cycle starts with an initial mean membrane potential that is higher than its asymptotic reference 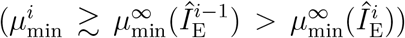, and hence has an instantaneous frequency that is (mostly) *higher* than the asymptotic reference (red color code). Vice versa, as the drive *decreases* (*m* < 0, Fig. 6B, right), trajectories will lie in the lower left half 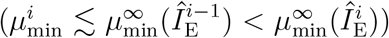 where instantaneous frequencies are mostly *lower* than their asymptotic reference (blue color code). Hence even a *symmetric*, piecewise linear double-ramp drive (Methods, Eq. (17)), induces the IFA asymmetry (Fig. 6C) : During the rising phase of the drive the instantaneous frequencies are above the asymptotic reference, and during the falling phase they lie below (Fig. 6C, top: markers vs black line; note cycle 5 as the only exception from the above “rule”). The IFA asymmetry thus does *not* rely on asymmetry in the input. Linear regression over the (semi-)analytically estimated instantaneous frequencies yields an IFA slope of −3.1 Hz/ms which is close to the spiking network simulation (Fig. 2B).

Interestingly, the last cycle *i* = 7 in Fig. 6C has a reference drive for which the constant-drive theory no longer applies 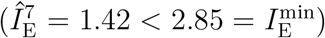, hence there is no asymptotic reference for the network frequency (empty marker, Fig. 6C, top). This means that at the end of the sharp wave the network can sustain one more ripple cycle at a level of drive that in the beginning would be insufficient to trigger ripples (see cycle 1 in Fig. 6C). The transient ripple is thus not only asymmetric in its instantaneous frequency (IFA), but also with respect to the level of drive at which it starts and ends.

We have established that IFA occurs in response to transient, sharp wave-like drive, independent of its symmetry, due to a hysteresis effect in the amplitude of the oscillatory mean membrane potential. Varying the slope *m* in our double-ramp model for SPW-like drive (Methods, Eq. (17)) we find that this hysteresis is *speed-dependent* (Fig. 7): If the drive changes more slowly, the transient dynamics approaches the asymptotic dynamics, and the IFA asymmetry is reduced (*χ*_IFA_ → 0 for |*m*| → 0, Fig. 7 A–C; Methods, Eqs. (58)-(68)). The theory thus explains the speed-dependence of IFA that we already observed in the spiking network simulations (Fig. 2D). The theoretically predicted instantaneous frequencies and IFA slopes are in excellent agreement with the corresponding observations in the spiking neural network simulation if the slope is sufficiently strong (Fig. 7 Aii-Bii, colored vs grey markers, average relative error *ϵ*: 14%, Table 1). The discrepancies between theory and simulation for slow drive (Fig. 7Cii, colored vs grey markers) are mainly due to the discrepancies in the estimate of the asymptotic reference frequencies for constant drive (Fig. 4).

**Table 1:** IFA in theory and simulation. Quantification of the IFA slope *χ*_IFA_ in the spiking network simulations and the theoretical approximations shown in Fig. 7Aii-Cii for different slopes *m* of the external SPW-like drive. The error *ϵ* (Eq. (69)) quantifies the average relative deviation of the theoretically predicted instantaneous network frequencies (colored markers in Fig. 7) from the simulation results (grey dots in Fig. 7).

| slope of SPW-like drive [1/ms] | $m = \pm 0.4$ | $m = \pm 0.2$ | $m = \pm 0.1$ |
| --- | --- | --- | --- |
| IFA slope $\chi_{\text{IFA}}$ (Hz/ms, simulation) | − 3.01 | − 0.68 | − 0.18 |
| IFA slope $\chi_{\text{IFA}}$ (Hz/ms, theory) | − 3.08 | − 1.75 | − 0.27 |
| error $\epsilon$ in instantaneous frequency | 14% | 14% | 13% |

**Figure 7:**
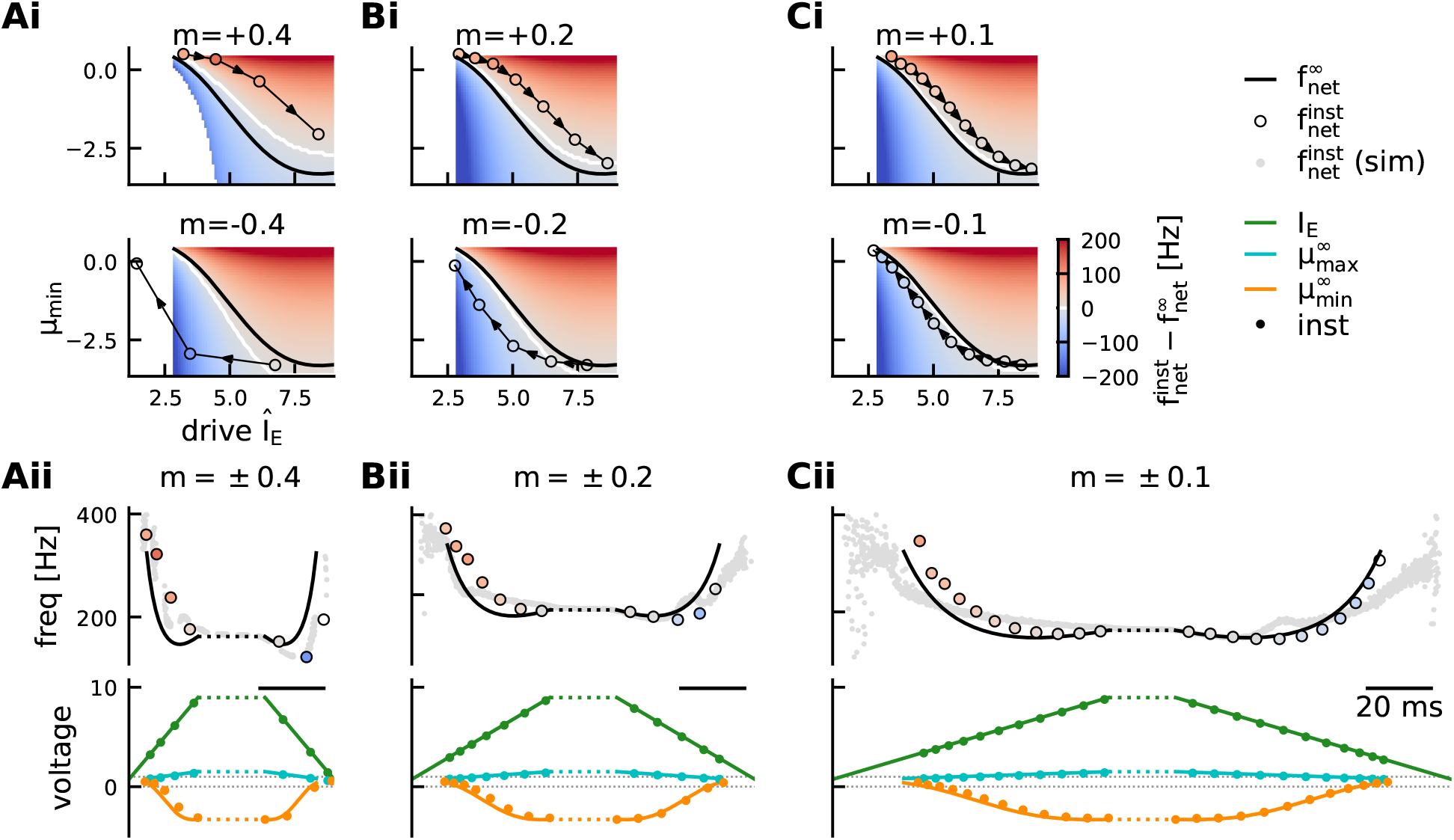
IFA is speed-dependent. Transient dynamics for SPW-like drive with slope *m* = ±0.4/ms (**A**, cf. Fig. 6), *m* = ±0.2/ms (**B**), or *m* = ±0.1/ms (**C**). Top panel **(i)**: difference between instantaneous and asymptotic network frequency for the possible combinations of external reference drive *Î*_E_ and initial mean membrane potential *μ*_min_, shown separately for positive (top) and negative (bottom) slope of the drive. Numerical trajectories shown in (ii) are overlaid. Bottom panel **(ii)**: Example trajectories through the space shown in (i) for SPW-like drive with slope ±*m*. Top: instantaneous (colored markers) vs asymptotic (black) network frequency as predicted by the theory. Grey dots indicate instantaneous frequencies in spiking network simulations (cf. Fig. 2D, *N* = 10, 000). See Table 1 for a quantitative comparison of simulation and theory. Bottom: transient vs asymptotic *μ*_min_ (orange) and *μ*_max_ (cyan). Green line shows SPW-like external drive (Eq. (17)), green dots mark reference drive *Î*_E_ of each cycle. The difference between instantaneous and asymptotic network frequencies (IFA) becomes less pronounced for smaller slope (C vs A, see also Table 1).

The speed-dependence of IFA is an important prediction of the bifurcation-based inhibitory ripple model that can be tested in experiments: Optogenetic stimulation of PV^+^ basket cells can trigger ripple oscillations (Schlingloff et al., 2014, but see Stark et al., 2014). Increasing and decreasing the intensity of the light pulse could mimick the piecewise linear, double-ramp drive studied here. The model predicts that the IFA asymmetry is reduced if the optogenetic light stimulus changes more slowly.

## Discussion

We have approximated the mean-field dynamics of the bifurcation-based inhibitory ripple model in the regime of strong drive and strong coupling. For constant drive, our theory (1) yields an approximation of the asymptotic network frequencies and mean unit firing rates far beyond the Hopf bifurcation, (2) captures the transition from sparse to full synchrony, and (3) reveals an increase of the mean membrane potential oscillation amplitude for increasing levels of external drive. For a fast changing, sharp-wave like drive we then show that a speed-dependent hysteresis effect in the trajectory of the mean membrane potential produces an IFA-like asymmetry of the instantaneous ripple frequency compared to the asymptotic frequencies. Our derivation shows that IFA is an intrinsic feature of the bifurcation-based inhibitory ripple model that will occur for any fast-enough sharp wave-like drive, independent of other parameter choices. The speed-dependence of IFA is a new prediction that can be tested experimentally.

To achieve an analytical treatment of the spiking network dynamics, we have made a number of simplifying assumptions. Our network dynamics are qualitatively similar to a biologically more realistic model (Donoso et al., 2018) that includes random sparse connectivity, correlations in the background noise, synaptic filtering, refractoriness, and conductance-based synapses.

Our reduced network model allows a Gaussian-drift approximation of the associated FPE dynamics that can be treated analytically. We added a phenomenological account for the reset mechanism on the population level, which improves the accuracy and range of applicability of the Gaussian-drift approximation. We would like to stress that this phenomenological reset is not *necessary* to capture the most important *qualitative* features of the network, namely the transition from sparse to full synchrony for constant drive and the hysteresis effect leading to IFA for time-dependent drive.

Our theory predicts that instantaneous ripple frequencies for a changing drive are different from the asymptotic frequencies for constant drive; for a double-ramp drive, the ripple frequency is higher during the rising and lower during the falling phase of the external drive. The concrete shape of the resulting intra-ripple frequency curve over time thus also depends on the shape of the asymptotic frequency as a function of the external drive. The asymptotic frequencies decrease for increasing values of the constant drive (Fig. 1B, in the vicinity of the Hopf bifurcation and in the limit of small synaptic delay this was already predicted by Brunel and Hakim (1999)). Thus it is possible for the instantaneous frequencies in the second half of the ripple event (when the external drive decreases) to lie *below* the asymptotic reference while actually *in*creasing as a function of time (Fig. 2D). Interestingly, such a switch from decreasing to increasing frequency at the end of the ripple event was also found in some experimental data (Ponomarenko et al., 2004). Others have reported an almost monotonic decrease or even a small peak in the beginning of the event (Nguyen et al., 2009; Sullivan et al., 2011; Stark et al., 2014; Donoso et al., 2018). For this reason we decided to put the emphasis on the overall asymmetry, rather than the exact shape of the instantaneous ripple frequency when studying IFA. Generally, intra-ripple frequency accommodation is robust with respect to the method used to estimate instantaneous frequencies (peak-to-peak distance vs. wavelet spectrogram, choice of time windows and smoothing kernels), even though there are differences, especially in the low power regime at the beginning and end of each event (Fig. 2A, discrete vs. continuous estimate).

We modeled the excitatory current input to CA1 during a sharp wave as a symmetric double ramp to highlight that IFA in this model does not depend on asymmetry in the input. Our derivation emphasizes that the only necessary requirement for the hysteresis causing IFA is an external drive that changes sufficiently fast and first rises, then decays. Thus we predict that IFA in the bifurcation-based model will occur for any such external drive, even when it is non-linear or asymmetric.

Other ripple models make different predictions w.r.t. IFA. In the “perturbation-based inhibitory model” (Malerba et al., 2016) the sharp wave-associated drive does not trigger a bifurcation but merely *perturbs* the network activity, which then settles into a new stable focus accompanied by transient oscillations, i.e. a ringing response. The network frequency reflects the average unit firing rate (no sparse synchrony) and thus depends directly on the strength of the external drive. A symmetric “up-down” drive thus creates a symmetric instantaneous frequency response that first rises and then decays. Since the oscillation power in this model decays monotonically over the course of the ripple event, the initial phase of rising frequency (“anti-IFA”) dominates over the subsequent decay. The perturbation-based model thus predicts that IFA only occurs in response to *asymmetric* drive (e.g. a sudden step up, followed by a ramp down). Excitatory currents measured during spontaneous SPW-Rs can exhibit some asymmetry due to synaptic filtering (Maier et al., 2011), but generally have a non-zero rise time (CA1 pyramidal cell: Maier et al., 2011; Donoso et al., 2018, CA3 PV^+^ BC: Hajos et al., 2013; Schlingloff et al., 2014). This speaks for the bifurcation-based inhibitory model, which can account for IFA being independent of the exact (a)symmetry of the SPW-associated drive. In both excitation-based ripple models, the ripple frequency depends on the average spike propagation delay among pyramidal cells — either orthodromically via supralinear dendrites (Memmesheimer, 2010; Jahnke et al., 2015) or antidromically via axo-axonal gap junctions (Traub et al., 1999; Traub and Bibbig, 2000). These models may be able to account for IFA by assuming an increase in the spike propagation delay over the course of a ripple event. Such an increase in latency might occur due to increasing somatic depolarization which has been shown to decrease the action potential amplitude (Shu et al., 2006, 2007). Future work should investigate in more depth whether and under which conditions excitation-first models can generate IFA.

Finally, understanding ripple oscillations requires not only understanding their generating mechanism but also the origin of the signal that we use to detect them. The local field potential is traditionally believed to reflect synaptic currents (Mitzdorf, 1985; Buzsáki et al., 2016), but modeling studies have suggested that action potentials of pyramidal cells can also contribute when occurring locked to a fast ripple rhythm (Schomburg et al., 2012; Ramirez-Villegas et al., 2018; Stark et al., 2014). Here we used the population activity of interneurons as a proxy for the LFP signal. It is reasonable to assume that postsynaptic inhibitory currents would exhibit the same frequency structure as the inhibitory population activity (Donoso et al., 2018). Simulations by Donoso et al. (2018) have shown that the inhibitory network can entrain the local pyramidal cell network, such that pyramidal cell spikes would occur phase-locked to the inhibitory ripple rhythm. A detailed model of the CA1 LFP would be needed to confirm whether any given generating mechanism yields a ripple signature in the LFP that is consistent with experimental data.

In conclusion: The inhibitory bifurcation-based ripple model can account naturally for IFA without adding further parameter constraints. A deepened understanding of the transient ripple dynamics in each of the proposed models together with extensive experimental testing of the various predictions will hopefully advance our understanding of the generating mechanism of ripple oscillations and enable us to study their potential role in memory consolidation.

## Materials and Methods

### Spiking neural network simulations

#### Network architecture

To model ripples in a spiking neural network, we consider a fully-connected inhibitory network of noisy leaky integrate-and-fire (LIF) neurons. Each neuron’s membrane potential *v_i_* (for *i* = 1, …, *N*) is given by the following stochastic differential equation (SDE):

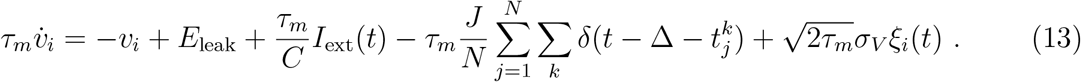

Whenever the membrane potential reaches a threshold of *V*_thr_ = −52 mV, a spike is recorded and the membrane potential is reset to *V*_reset_ = −65 mV. For simplicity there is no absolute refractory period. We choose a membrane time constant *τ_m_* = 10 ms, resting membrane potential *E*_leak_ = −65 mV and membrane capacitance *C* = 100 pF as used by Donoso et al. (2018). All default parameter values are summarized in Table 2.

**Table 2:** Default parameters for the spiking network

| Parameter | Value | Definition |
| --- | --- | --- |
| $N$ | 10,000 | Number of interneurons |
| $\tau_m$ | 10 ms | Membrane time constant |
| $C$ | 100 pF | Membrane capacitance |
| $E_{\text{leak}}$ | -65mV | Resting potential |
| $V_{\text{thr}}$ | -52 mV | Spike threshold |
| $V_{\text{reset}}$ | - 65 mV | Reset potential |
| $J$ | 65 mV | Inhibitory coupling strength |
| $\Delta$ | 1.2 ms | Synaptic delay |
| $\sigma_V$ | 2.62 mV | Standard deviation of Gaussian white noise input |
| $V_T$ | 1 | Spike threshold, dimensionless |
| $V_R$ | 0 | Reset potential, dimensionless |
| $K$ | 5 | Inhibitory coupling strength, dimensionless |
| $D$ | 0.04 | Variance of Gaussian white noise input, dimensionless |

In our model, Eq. (13), each unit receives a common excitatory drive *I*_ext_(*t*) and a common inhibitory synaptic input in form of a sum of presynaptic Dirac-delta spikes from all neurons in the network. Spikes occur at times 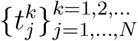, where 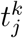 denotes the *k*-th spike time of neuron *j*. They are transmitted to all neurons in the network and arrive at the postsynaptic neurons after a synaptic delay of Δ = 1.2 ms. The amplitude of the inhibitory postsynaptic potential elicited by one input spike is determined by the synaptic strength *J* = 65 mV. Our default network has *N* = 10, 000 units.

To account for noisy background activity, every unit receives independent Gaussian white noise *ξ_i_*(*t*) with ⟨*ξ_i_*(*t*)⟩ = 0, ⟨*ξ_i_*(*t*)*ξ_j_*(*t*′) = *δ_i,j_δ*(*t* − *t*′). The strength of the noise is determined by the parameter *σ_V_* = 2.62 mV, which can be interpreted as the long-time standard deviation of the membrane potential in the absence of a threshold.

In simulations of the spiking network with a finite temporal resolution (Δ*t* = 0.01 ms) the *empirical* population activity is estimated as

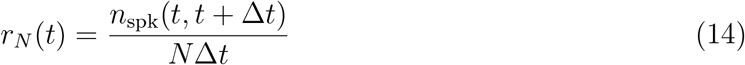

where

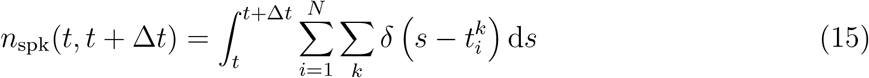

denotes the total number of spikes that were emitted from the population in the time interval [*t, t* + Δ*t*]. *r_N_* has units of spikes per second and can also be interpreted as the instantaneous firing rate of any given neuron in our homogeneous network (Gerstner and Kistler, 2002).

For plotting purposes the empirical population activity is smoothed with a Gaussian window *g* of standard deviation *σ_t_* = 0.3 ms:

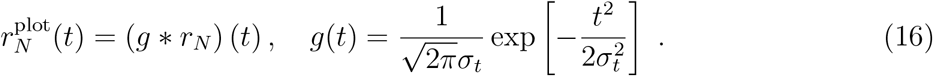

To facilitate notation we omit the superscript “plot” in all Figures.

For constant drive, we simulate 5.05 s (time step Δ*t* = 0.01 ms). The initial 50 ms are excluded from analysis, because we are not interested in the initial transient but in the asymptotic network dynamics. The remaining 5 s are sufficient for a basic spectral analysis.

For time-dependent drive, we first simulate the network for 200 ms with a constant baseline drive of 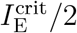 (see “Linear stability analysis”), followed by the time-dependent, *sharp wave-like* stimulus, which we model as a piecewise linear double-ramp of slope ±*m* with a plateau phase of arbitrary length in between (here 20 ms):

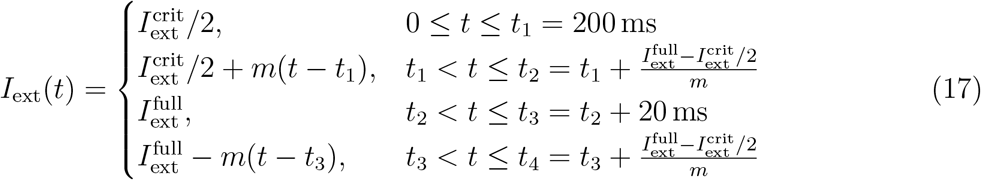

During the plateau phase the drive is at the approximate point of full synchrony 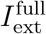 for the respective spiking network, as determined by a range of constant-drive simulations.

Simulations of the LIF network model have been performed using the Brian2 simulator (Goodman and Brette, 2009). Data storage and parallelization of simulations for large parameter explorations were done using the Python toolkit *pypet* (Meyer and Obermayer, 2016).

#### Frequency analysis

The network frequency at constant drive is defined as the location of the dominant peak in the power spectral density of the population activity *r_N_*(*t*). The saturation (average fraction of neurons firing in one cycle of the population rhythm) is computed by dividing the mean unit firing rate by the network frequency.

To define the instantaneous network frequency in response to time-dependent drive, we use frequency estimates both in continuous time and for a discrete set of time points. The continuous estimate is derived from the wavelet spectrogram (windowed Fourier transform) of the population activity, which indicates instantaneous power in the frequency band from 0 to 350 Hz over time; the instantaneous frequency at each point in time is defined as the frequency above 70 Hz, that has maximal instantaneous power. The lower limit is introduced to exclude the low-frequency contribution due to the sharp wave. The instantaneous frequency is considered significant, whenever the corresponding instantaneous power exceeds a power threshold. The power threshold is chosen as the average instantaneous power at 0 Hz during the initial 200 ms baseline window, plus 4 standard deviations.

An alternative estimate of the instantaneous frequency is given by the inverse of the peak-to-peak distances in the (smoothed) population rate. We consider only peaks that are more than 4 standard deviations above the average rate during baseline stimulation. This procedure delivers a discrete set of frequency estimates associated with a discrete set of time points. In this paper we mostly rely on this “discrete” estimate of instantaneous frequency since its parameter-dependencies (minimal height of oscillation peaks) are more transparent than the ones of the continuous-time estimate (size of time window for windowed Fourier transform, power threshold). Furthermore it is better suited for comparison with the theory which also describes instantaneous frequency as a discrete measure per cycle (see Eq. (67)).

To quantify the network’s instantaneous frequency response to a SPW-like drive (Eq. (17)), we perform 50 independent simulations of the network model with the same drive but different noise realizations. Linear regression over the discrete instantaneous frequency estimates 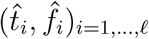 from *ℓ* discrete oscillation cycles, pooled together from all 50 simulations, yields the average change of the instantaneous frequency over time:

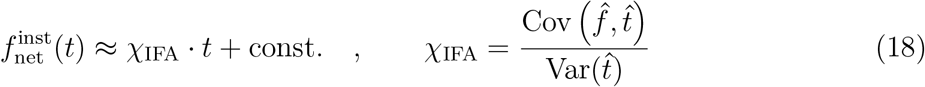

A negative slope *χ*_IFA_ < 0 indicates IFA.

Note that for our symmetric model of SPW-like drive (Eq. (17)), the regression slope quantifies the change in the *deviations* of the instantaneous network frequencies 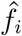 from the symmetric, asymptotic reference frequencies 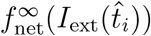 (except for small effects of asymmetric sampling):

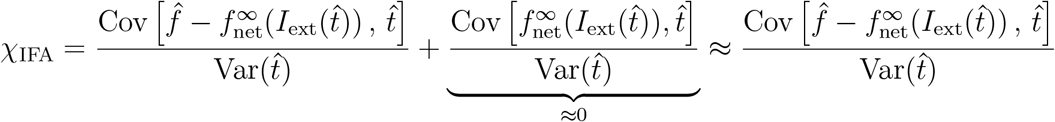

#### The point of full synchrony

We estimate the point of full synchrony in the spiking network by interpolating the simulated saturation curve and estimating the level of drive for which it becomes 1.

#### Extracting the average oscillation cycle

For constant drive, we split the spiking network simulation into individual cycles based on the Hilbert transform of the mean membrane potential. We take a sufficient number of equally spaced samples from each cycle (here 21) and average them across cycles to derive the average trajectory of the population rate and the membrane potential histogram over the course of a ripple cycle. For each of the 21 sample times we can calculate the average membrane potential, which will be used for comparison with the theory.

### Mean-field approximation

#### Dimensionless equations

To facilitate notation in the theoretical part of this paper, we shift and rescale all voltages, such that the spiking threshold becomes *V_T_* = 1 and the resting potential is *E_L_* = 0. This corresponds to measuring voltage in units of the distance from the resting potential to the spike threshold. The single unit SDE then reads

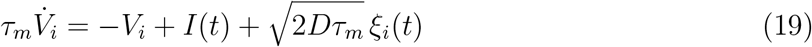

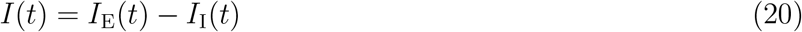

where now

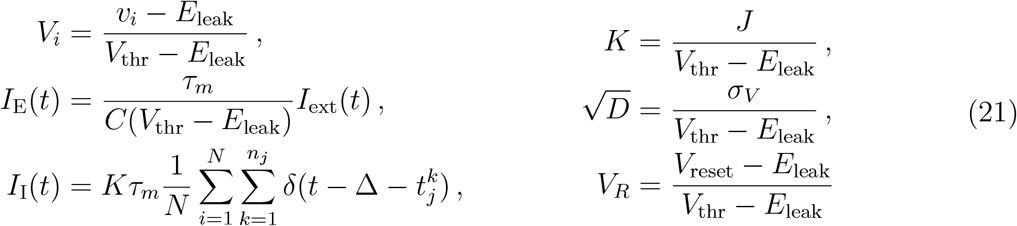

are all dimensionless quantities.

#### Fokker-Planck equation

In the mean-field limit of an infinitely large interneuron population (*N* → ∞) the evolution of the membrane potential density *p*(*V, t*) is described by the following Fokker-Planck equation (FPE) (see e.g. Abbott and van Vreeswijk, 1993; Brunel and Hakim, 1999):

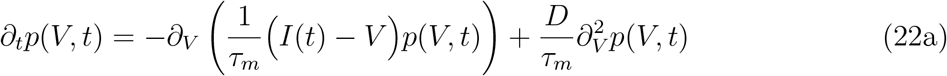

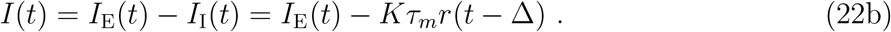

The FPE can also be written as a continuity equation

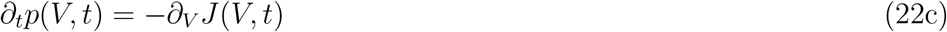

with a probability current

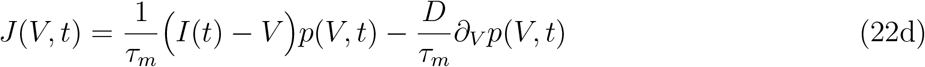

Since units are reset instantaneously as soon as they reach the spiking threshold, the FPE has an absorbing boundary condition at threshold:

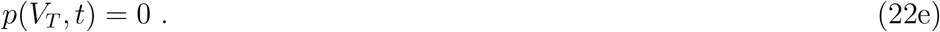

The population rate *r* is given by the probability current through the threshold:

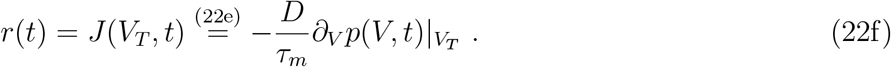

The fire-and-reset mechanism introduces a derivative discontinuity at the reset potential *V_R_*:

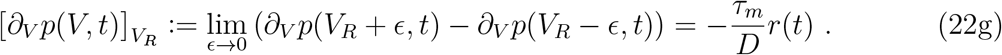

(Abbott and van Vreeswijk, 1993; Brunel, 2000; Lindner and Schimansky-Geier, 2001; Fourcaud and Brunel, 2002; Brunel et al., 2003). The membrane potential density must decay to zero fast enough in the limit of *V* → −∞:

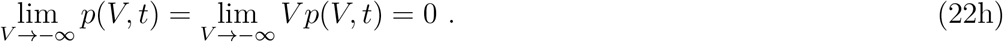

As a probability density, *p*(*V, t*) obeys the normalization condition:

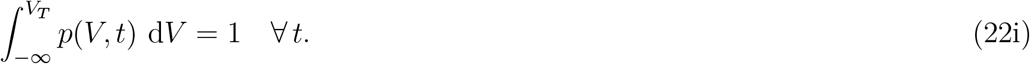

#### Stationary solution

The stationary solution *p*_0_(*V*) of the FPE has been derived by Brunel and Hakim (1999) (see also Abbott and van Vreeswijk, 1993). The constant population rate *r*_0_ and resulting total drive *I*_0_ in the stationary state can be inferred self-consistently by solving:

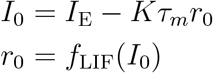

where

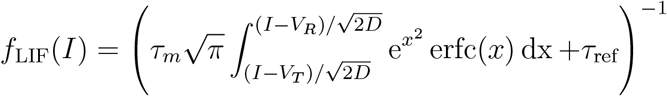

is the firing rate of an uncoupled LIF neuron receiving constant drive *I* and Gaussian white noise of intensity *D* (f-I curve, Holden, 1976).

#### Linear stability analysis

For a given external drive *I*_E_ we assume a weak, periodic perturbation of the population rate around its stationary value:

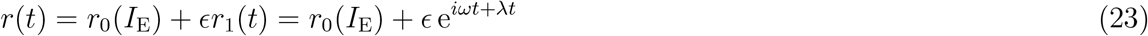

The bifurcation where the stationary state loses stability corresponds to *λ* = 0. In the recurrently coupled network this perturbation of the rate translates into a perturbation of the input current:

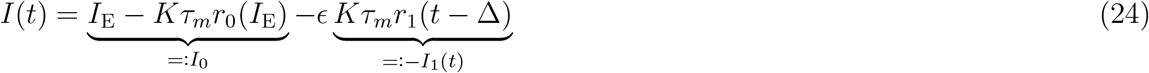

The linear response of the LIF units to this weakly modulated drive *I*(*t*) under Gaussian white noise is given by convolution with the linear response function *G*:

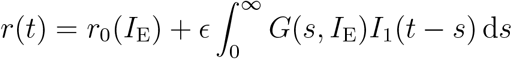

This output rate must match the weakly periodically modulated rate *r*(*t*) that we assumed in the beginning (Eq. (23)):

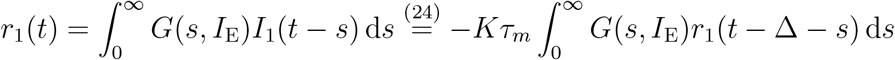

At the bifurcation (*r*_1_(*t*) = e^*iωt*^, Eq. (23)) this self-consistent condition is equivalent to

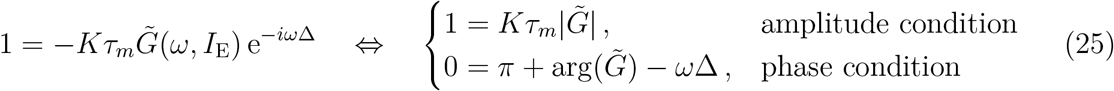

where 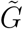 denotes the Fourier transform of the linear response function (*susceptibility*). We use the exact expression for the susceptibility of an LIF unit under Gaussian white noise (more specifically, the complex-conjugated of the expression derived by Lindner and Schimansky-Geier (2001)):

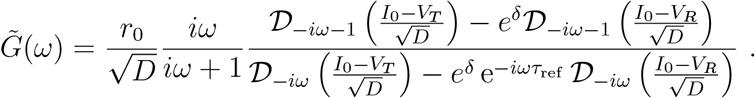

where 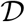 are parabolic cylinder functions and 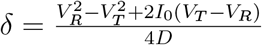 (for an alternative expression in terms of confluent hypergeometric functions, see Brunel et al., 2001). We solve the amplitude and phase condition (Eq. (25)) numerically to find the critical drive 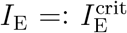 (at which the stationary state loses stability) and the corresponding frequency *ω* of the emerging oscillation. The network frequency and mean unit firing rate at the bifurcation are thus given by 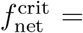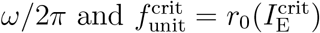. This approach is equivalent to the derivation by Brunel and Hakim (1999) via linear expansion of the FPE solution (see also Brunel and Hansel, 2006).

### Derivation of the Gaussian-drift approximation

Without the absorbing boundary at threshold and the source term due to the fire-and-reset rule, the FPE has only natural boundary conditions and can be solved analytically. Its solution *p_δ_* for an initial Dirac delta distribution *p_δ_*(*V*, 0) = *δ*(*V* − *μ*_0_), is a Gaussian density with time-dependent mean *μ*(*t*) and variance *σ*^2^(*t*) (Uhlenbeck and Ornstein, 1930):

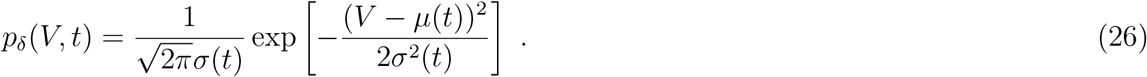

The mean membrane potential *μ* evolves according to the single unit ODE (Eq. (19)) without the noise term:

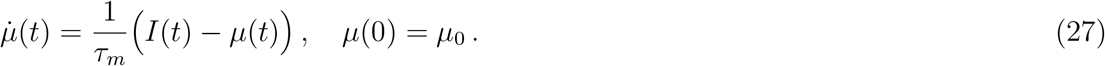

The variance of the membrane potential distribution, *σ*(*t*)^2^, approaches *D* with a time constant of *τ_m_*/2:

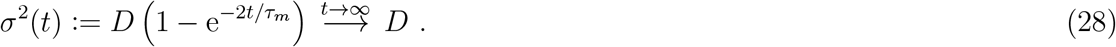

The solution for an arbitrary initial condition can be found by convolution of the initial condition with *p_δ_*. Hence in the long time limit (*t* → ∞) *all* solutions of the FPE with natural boundary conditions tend towards a Gaussian with variance *D* — independent of the initial condition (which has to satisfy the boundary conditions Eqs. (22)).

In our ripple-generating network, with strong drive and strong coupling, the bulk of the membrane potential distribution is strongly *sub*threshold in between population spikes (*i.e*. unaffected by the non-natural boundary conditions), and thus tends towards a Gaussian density. When the next population spike begins, we can hence approximate the membrane potential density as a Gaussian with fixed variance *D*:

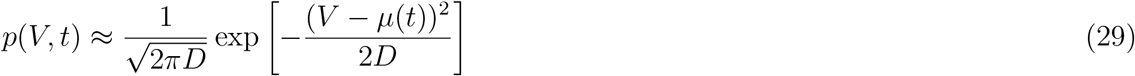

This Gaussian approximation allows the derivation of a simple expression for the population rate under strong drive (Plesser and Gerstner, 2000; Chizhov and Graham, 2007; Goedeke and Diesmann, 2008):

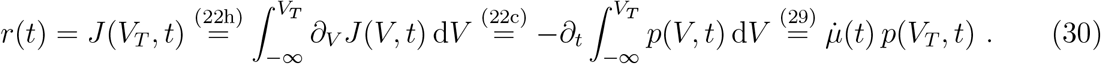

The rate is given by the value of the Gaussian density at threshold, scaled by the speed at which the mean membrane potential approaches the threshold. Since only *upwards*-threshold-crossings should contribute to the rate, we add a sign-dependence:

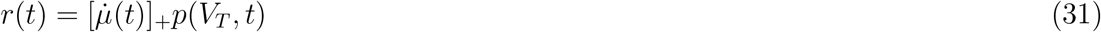

clipping the rate to 0 whenever the mean membrane potential decays: 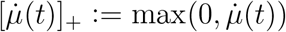 (Plesser and Gerstner, 2000; Chizhov and Graham, 2007; Goedeke and Diesmann, 2008).

#### Numerical analysis of oscillation dynamics

In its derivation above, we formulated the Gaussian-drift approximation in several equations, describing the membrane potential density *p* (Eq. (29)), mean membrane potential *μ* (Eq. (27)) and population rate *r* (Eq. (31)) separately. Note however that the mean membrane potential *μ* is the only independent variable and thus the Gaussian-drift approximation can be rephrased as a single delay differential equation (DDE):

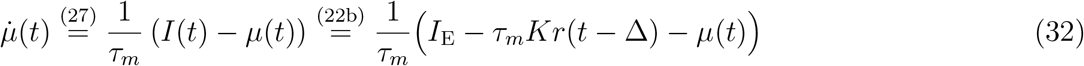

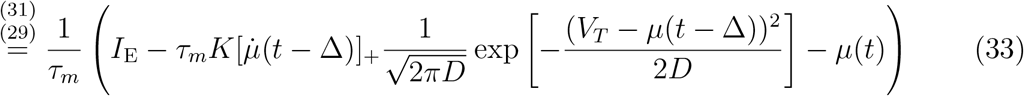

We will nevertheless illustrate the solutions as two-dimensional trajectories of both *μ*(*t*) and *r*(*t*) (Fig. 8), since the population rate is our main variable of interest. Using a simple forward Euler method and initial condition *μ*(0) ≪ *V_T_* such that *r*(*t*) ≈ 0 ∀ *t* ≤ 0, we can numerically integrate the DDE and find a range of potential dynamics for constant drive *I*_E_ (see Fig. 8).

**Figure 8:**
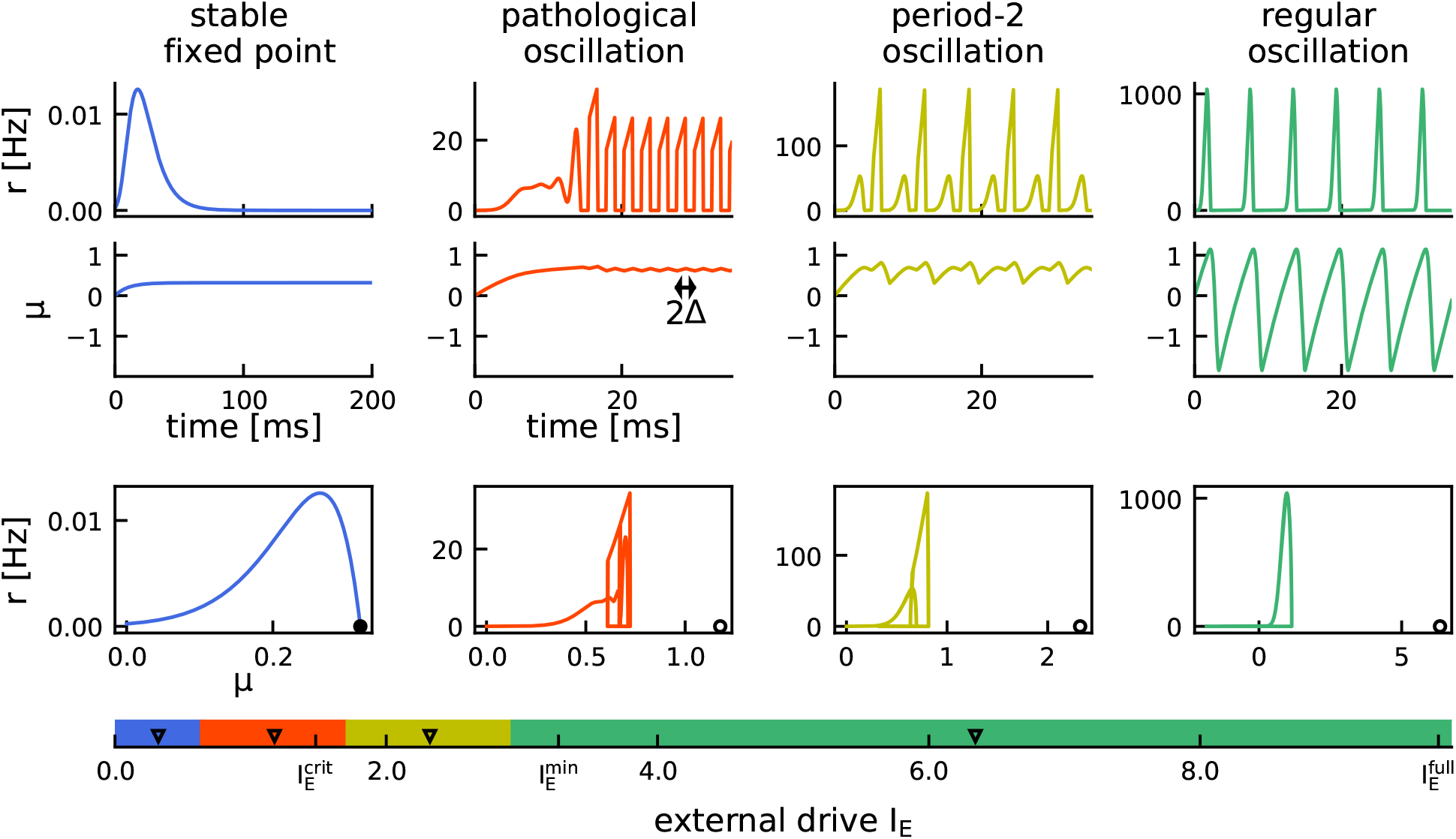
Dynamical states of the DDE system. Numerical integration of the DDE system demonstrating 4 distinct dynamical regimes for increasing external drive *I*_E_ (bottom axis). Blue: Stable fixed point with zero rate. Red: Pathological, fast oscillations. Yellow: Period-2 oscillations. Green: Regular period-1 oscillations. Top: Numerically integrated trajectories of population rate *r* and mean membrane potential *μ* over time. Note the changes in scale for the population rate. Bottom: Phase space showing the trajectory (*μ*(*t*), *r*(*t*)) and the fixed point (*I*_E_, 0), which is only stable in the first case (black circle) and unstable otherwise (empty circle). The horizontal colored bar shows the full range of (relevant) drives from 0 to the point of full synchrony 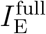. Black triangles mark the levels, for which the above example dynamics are shown. Extra ticks indicate: critical drive 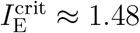, for which the spiking network undergoes a Hopf bifurcation (see “Linear Stability Analysis”); lower bound 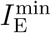 introduced for our theory to ensure that we only consider non-pathological dynamics (see “Range of applicability”).

There is a large regime of sufficiently strong drive, in which the solution *μ* exhibits persistent period-1 oscillations (Fig. 8, green). This is the regime that we are interested in and the dynamics of which we will approximate in the following.

At lower levels of drive there are three additional dynamical regimes that we will exclude from analysis: At low drive, the system has a stable fixed point 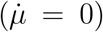 in (*μ*(*t*), *r*(*t*)) ≡ (*I*_E_, 0) (Eq. (32), Fig. 8, blue). The bifurcation at which the fixed point loses stability can be determined numerically. Immediately after the bifurcation the DDE solution exhibits very fast oscillations at (2Δ)^−1^ ∼ 417 Hz (Fig. 8, red). The oscillation amplitude of the mean membrane potential is very small and a large portion of the Gaussian potential density is suprathreshold at all times. We refer to this oscillation as pathological since it is a direct result of the artificial clipping of the rate to 0 whenever the mean membrane potential decays (Eq. (31)). Increasing the drive further brings the system into a state of period-2 oscillations where the Gaussian density gets pushed below threshold only every other cycle (Fig. 8, yellow).

These regimes of pathological high frequency or period-2 oscillations exist due to our simplifying assumptions capturing only the mean-driven aspects of the network dynamics. Both regimes occur either shortly before or after the level of drive 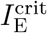 at which the original spiking network undergoes a supercritical Hopf bifurcation (see tick mark in Fig. 8). In the vicinity of that bifurcation the spiking network dynamics are fluctuation-driven with either no 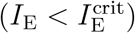 or only small-amplitude 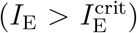 oscillations in the mean membrane potential and population rate. These cannot be captured without taking into account the absorbing boundary at threshold and the non-Gaussian shape of the density of membrane potentials. We will exclude these pathological dynamics from analysis by introducing a lower bound 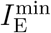 for the theoretical approximations that we develop in the following (see tick mark in Fig. 8).

### Analytical approximation of oscillation dynamics for constant drive

For strong enough drive the mean membrane potential *μ* under the Gaussian-drift approximation oscillates periodically between two local extrema *μ*_min_ and *μ*_max_ (Fig. 8, green regime, see also Fig. 9A). The population rate *r*(*t*) oscillates at the same frequency and is positive when the mean membrane potential increases, *i.e.* 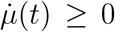, and 0 otherwise, c.f. Eq. (31). The time when the mean membrane potential reaches its local maximum *μ*_max_ marks the end of the population spike and will be denoted as *t*_off_. The inhibitory feedback (Eq. (22b)) thus ends at time *t*_off_ + Δ and we define *μ*_min_ ≔ *μ*(*t*_off_ + Δ) as the end of a cycle. This allows us to approximate the overall period as *T* = *t*_off_(*μ*_min_, *μ*_max_) + Δ. Here, we wrote *t*_off_(*μ*_min_, *μ*_max_) to emphasize that *t*_off_ is the rise time of the mean membrane potential from *μ*_min_ to *μ*_max_ (*up-stroke*). Furthermore, Δ is the duration of the subsequent *downstroke* back to *μ*_min_. In the following, we will approximate *μ*_max_ (Step 1) and *μ*_min_ (Step 2) and thus derive the network frequency as the inverse of the period:

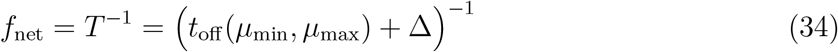

**Figure 9:**
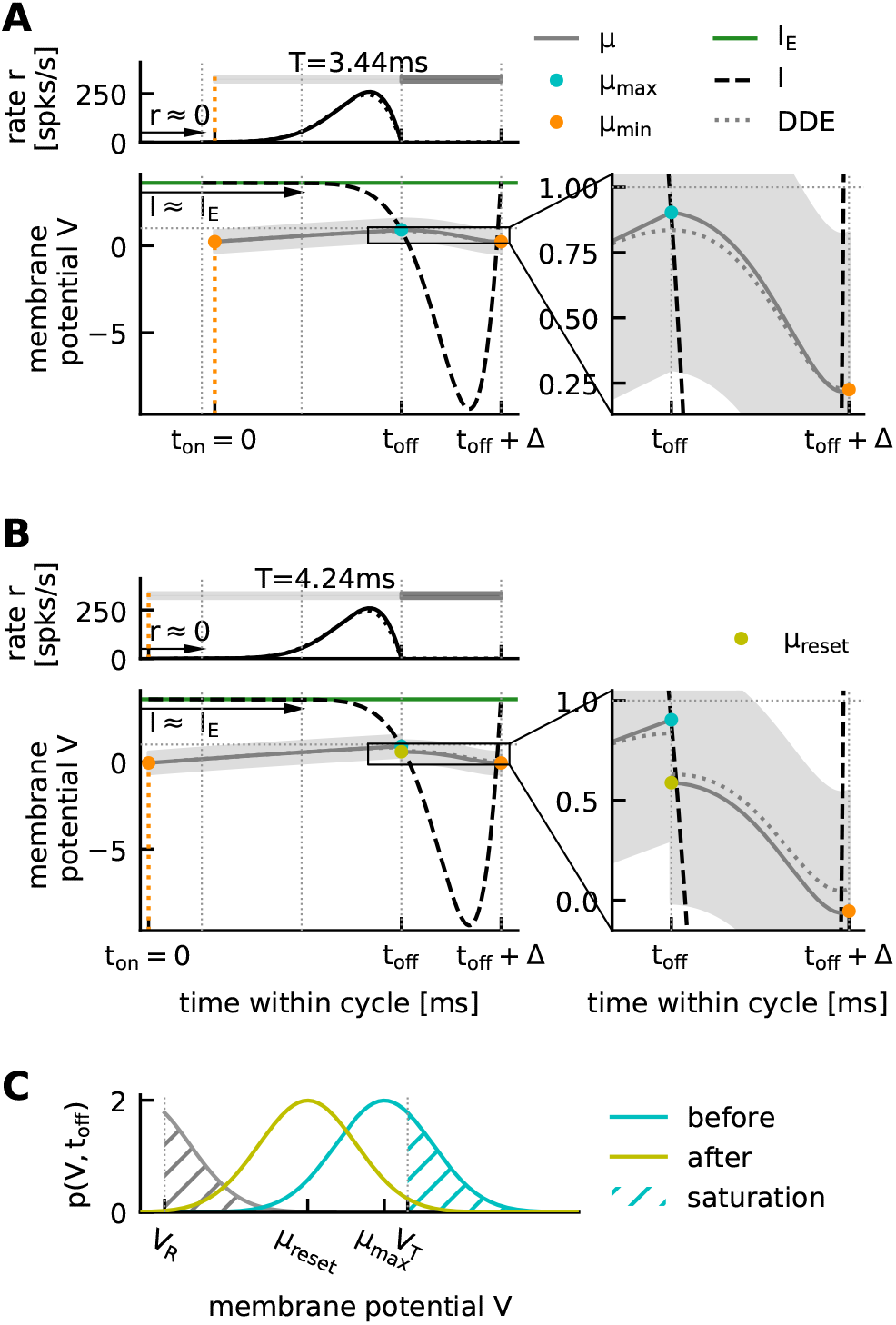
Illustration of analytical approximation of DDE dynamics. **A**, One cycle of the oscillatory solution of the DDE for *I*_E_ = 3.6. Dotted lines for rate (top) and mean membrane potential (bottom) are the result of a numerical integration of the DDE (Eq. (33)). All other lines illustrate our analytical considerations. Top: population rate *r*(*t*) (black). Bottom: mean membrane potential *μ*(*t*) (full grey line: Eq. (A2) during upstroke, Eq. (41) during downstroke, gray area: 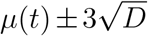 indicating the width of the Gaussian density *p*(*V, t*)). Constant external drive *I*_E_ (green line), total input *I*(*t*) = *I*_E_ – *I*_I_(*t*) (dashed line, Eq. (44)). Local extrema of the mean membrane potential occur at the intersections of *μ* and *I*: Cyan: local maximum *μ*_max_ (Eq. (40)). Orange: approximate local minimum *μ*_min_ (Eq. (45)). Vertical dotted lines mark end of population spike *t*_off_ as well as intervals *t*_off_ + *k*Δ, *k* ∈ {−2, −1, 1}. Arrows illustrate simplifying assumption (A1). The beginning of the cycle (*t*_on_ = 0) is determined by *μ*_min_. Horizontal gray bars mark the length of one cycle (here *T* = *t*_off_ + Δ = 3.44 ms (Eq. (49)), corresponding to a network frequency of *f*_net_ = 290.7 Hz, (Eq. (34))). Inset: magnification highlighting the differences between numerical solution (dotted) and analytical approximation (full line). Due to assumption (A2) *μ*_max_ is slightly overestimated. Note that the second intersection of *μ* and *I* occurs shortly *before* time *t*_off_ +Δ. Hence *μ*_min_ is slightly larger than the true local minimum. **B**, Same as A, but with an account for the reset on the population level. At the end of the population spike, *μ* is reset instantaneously from *μ*_max_ to *μ*_reset_ (Eq. (51)) (yellow marker). This leads to a lower *μ*_min_ (Eq. (52)) and hence a slightly longer period (*T* = 4.24 ms), *i.e*. lower network frequency (*f*_net_ = 235.8 Hz). **C**, Illustration of phenomenological reset. Cyan: density of membrane potentials *p*(*V, t*_off_) at the end of the population spike, centered at *μ*_max_ (before reset). Cyan hatched area: fraction of active units (*saturation*, Eq. (50)). Grey hatched area: resetting the active portion of *p*. Yellow: *p*(*V, t*_off_) after reset, centered at *μ*_reset_. The reset value *μ*_reset_ is calculated as the average of the density that results from summing the grey-dashed (active units) and cyan-non-hatched (silent units) density portions (Eq. (51)). Default parameters (see Table 2).

#### Step 1

To find the local maximum *μ*_max_ = *μ*(*t*_off_) we have to solve

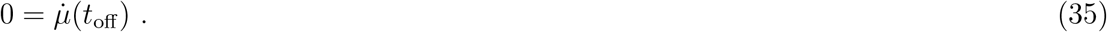

Since the dynamics of the mean membrane potential are given by a delay differential equation, the term on the right-hand side is recurrent:

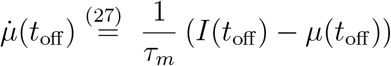

From the yet unknown time *t*_off_ we have to look back in time (in windows with length of the delay Δ) at the history of the population rate *r*(*t*):

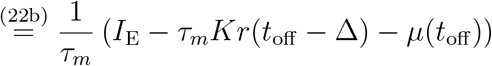

which in turn depends on *μ*(*t*) and 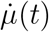:

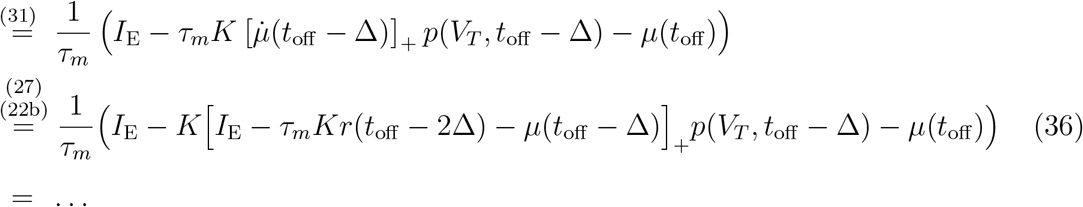

We can resolve the recurrence since there is only a *finite* number of past time windows, during which the population rate, and thus the delayed feedback inhibition *I*_I_(*t*) = *Kτ_m_r*(*t* – Δ), is significantly above 0. In the first time window [*t*_off_ – Δ, *t*_off_], right before the end of the upstroke, we have to take inhibition into account, since this is what stops the upstroke. In the second time window [*t*_off_ – 2Δ, *t*_off_ – Δ], further into the past, we will assume that the inhibitory feedback is negligible, *i.e*.

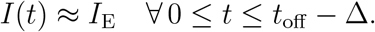

This assumption implies that the population rate was negligible in the previous time window, *i.e*.

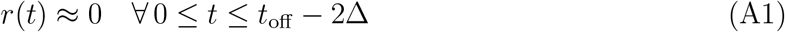

(Eq. (22b), Fig. 9A). Note that *t* here refers to time since the beginning of the cycle (*t* = 0). Since the population spike ends at time *t*_off_, (A1) is equivalent to the assumption that the population spike lasts at most 2Δ. Adding the subsequent downstroke time of Δ this amounts to an upper bound for the oscillation period of around 3Δ, plus any additional upstroke time with *r* ≈ 0, which is a reasonable assumption for a feedback loop with delay Δ as argued already by Brunel and Hakim (1999).

Under this assumption, we set *r*(*t*_off_ – 2Δ) = 0 in Eq. (36), and only a finite number of terms remains:

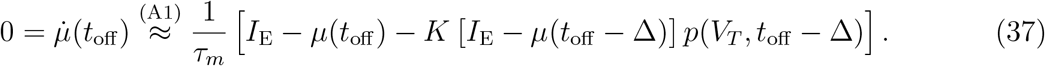

Note that the rectification [·]_+_ has been dropped in this equation because the mean membrane potential can never be larger than the external drive. Furthermore, we assume that we can approximate the past *μ*(*t*_off_ − Δ) during the upstroke by only considering the excitatory drive, *I*(*t*) ≈ *I*_E_. Under this assumption, Eq. (27) yields the exponential relaxation

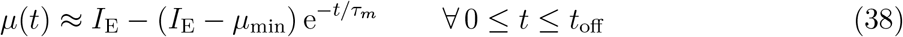

Looking backwards from time *t*_off_, we can reformulate this assumption as:

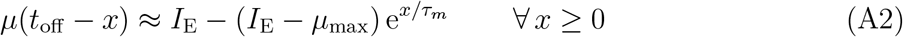

The resulting error is small since the exponential relaxation of the mean membrane potential towards the total drive *I*(*t*) is governed by the membrane time constant (*τ_m_* = 10 ms). The population spike on the other hand is quite synchronized (we assumed that it lasts less than 2Δ ≪ *τ_m_*, (A1)). The time window right before *t*_off_, during which the units receive inhibitory feedback and the total drive deviates from *I*_E_, is thus small compared to the membrane time constant and alters the trajectory of *μ* only slightly (Fig. 9A, inset). Equation (37) thus simplifies to

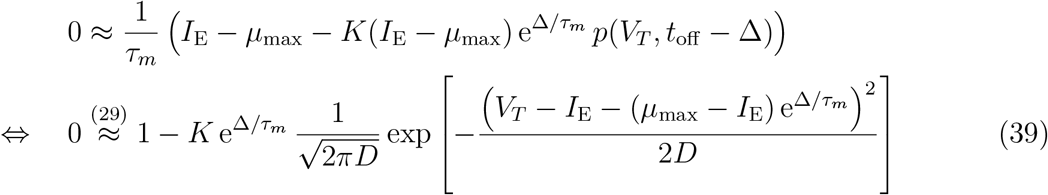

The simplified equation can be readily solved for *μ*_max_:

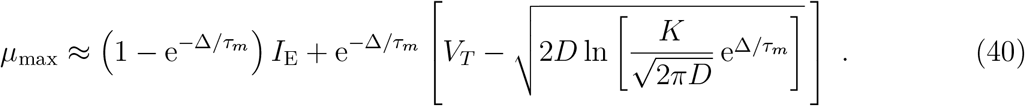

Note that *μ*_max_ does *not* depend on the initial voltage *μ*_min_. This makes sense intuitively for any *μ*_min_ that is sufficiently smaller than *μ*_max_, since the “turning point” of the mean membrane potential depends only on the feedback inhibition resulting from the immediate history shortly before is reached (*t* ∈ [*t*_off_ – 2Δ, *t*_off_]). To ensure that the argument of the logarithm in Eq. (40) is larger than 1, the coupling must be sufficiently strong: 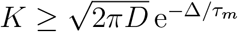.

#### Step 2

We will now approximate the trajectory of the mean membrane potential during its inhibition-induced downstroke and infer *μ*_min_ = *μ*(*t*_off_ + Δ), which corresponds to both the end and the starting value of each cycle of a periodic solution.

Note that while *μ*_min_ is close to the periodic local minimum of the mean membrane potential, it is a slight overestimation: The local extrema of the mean membrane potential *μ* occur at its intersections with the total drive *I*(*t*) (see Eq. (35), Fig. 9A). At time *t*_off_ the mean membrane potential reaches its local maximum and becomes larger than the total drive *I*(*t*) (Fig. 9A). Since the population spike ends at time *t*_off_, the delayed inhibitory feedback *I*_I_(*t*) will stop at time *t*_off_ + Δ: The total drive at this point will equal the external drive (*I*(*t*_off_ + Δ) = *I*_E_); note that the mean membrane potential *μ* can never be larger than the external drive *I*_E_. Hence, if *μ* becomes larger than *I* at time *t*_off_ and is smaller than *I* at time *t*_off_ + Δ, *μ* must intersect with *I*(*t*) and reach its local minimum slightly *before* time *t*_off_ + Δ (see Fig. 9A, inset). What we define as the initial/final membrane potential of a cycle (*μ*_min_ ≔ *μ*(*t*_off_ +Δ)) is thus close to but slightly larger than the periodic local minimum. This does not affect our estimate of the period.

The definition of a fixed downstroke duration Δ allows us to find *μ*_min_ directly by integrating the mean membrane potential ODE Eq. (27) up to time *t*_off_ + Δ, starting from the initial value *μ*(*t*_off_) = *μ*_max_:

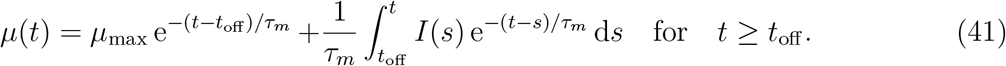

It follows for *μ*_min_ = *μ*(*t*_off_ + Δ) with initial condition *μ*(*t*_off_) = *μ*_max_:

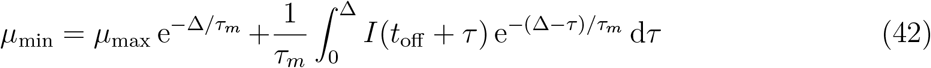

Again, the total current *I*(*t*_off_ + *τ*) with *τ* ∈ [0, Δ], has a recurrent dependency on the past dynamics, which can be seen by repeated application of Eqs. (22b),(31),(27):

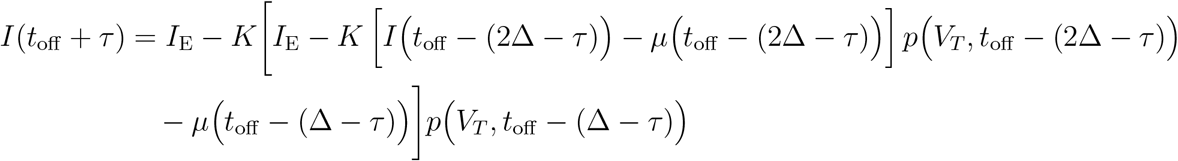

As before we assume that the current *I* at time *t*_off_ (2Δ – *τ*) ≤ *t*_off_ − Δ is given exclusively by the excitatory drive (A1), which truncates the infinite recurrent expression above to:

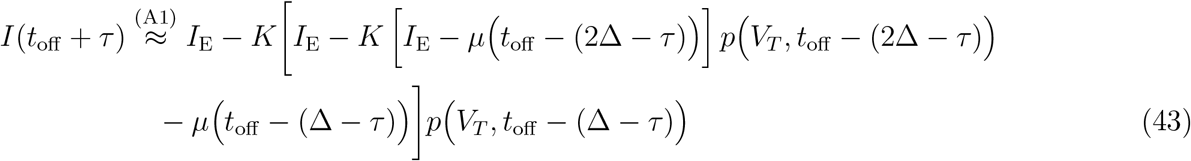

We are left with a finite number of terms that depend on the trajectory of *μ* (*t*) during the upstroke. Again, we approximate *μ*(*t*_off_ – *x*), *x* ≥ 0 assuming exponential relaxation towards only the external drive, *i.e*. ignoring inhibition (A2). This abolishes all dependencies on the yet unknown time *t*_off_ of the end of the population spike:

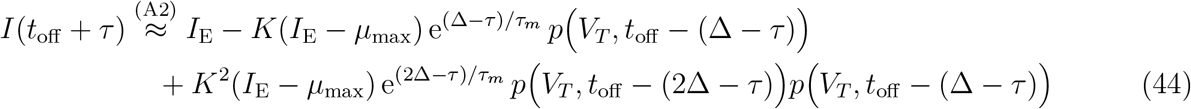

Inserting this approximation (Eq. (44)) into the expression for *μ*_min_ (Eq. (42)) yields:

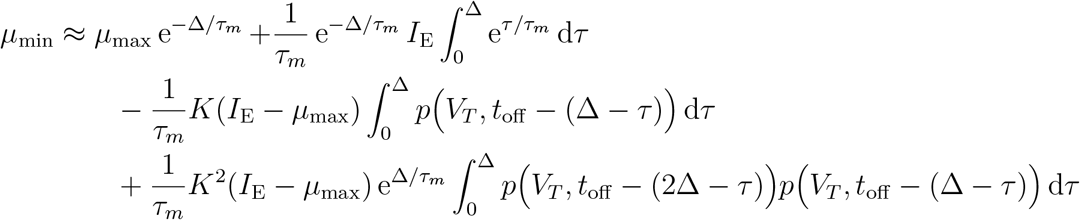

The integrals can be solved analytically, if we approximate the past trajectory of *μ* linearly within the Gaussian expressions *p*:

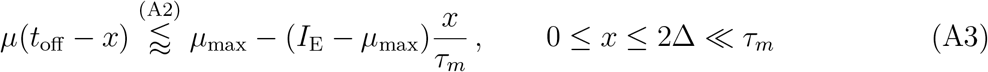

Hence,

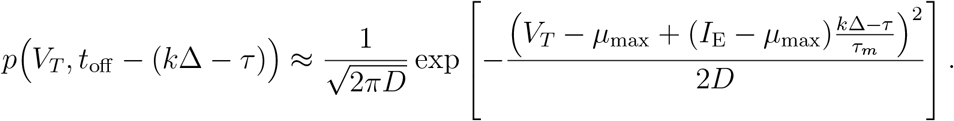

Using this expression, we obtain

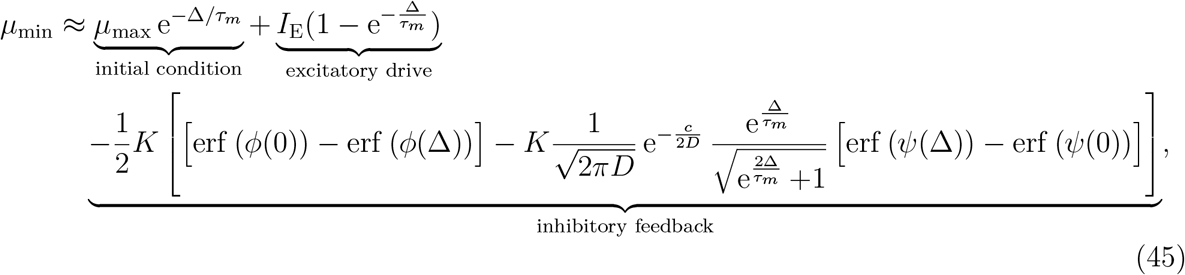

where

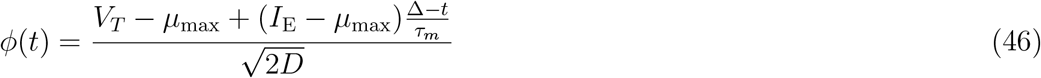

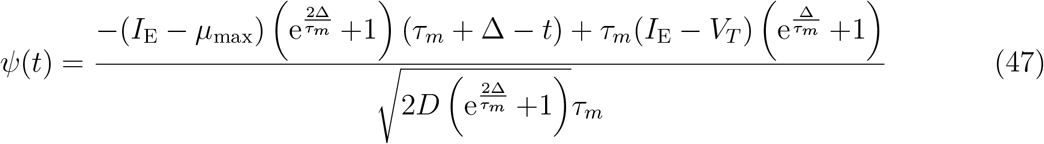

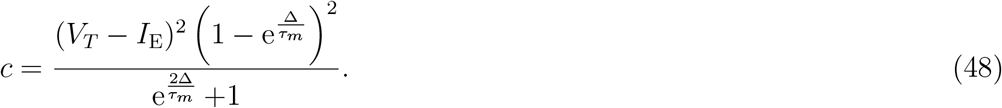

Although lengthy, this expression can be easily evaluated numerically.

The local “minimum” *μ*_min_ is not only the final but also the initial mean membrane potential of each cycle. The length of the upstroke can thus be approximated as the time it takes for the mean membrane potential to rise from *μ*_min_ to *μ*_max_, based on exponential relaxation towards only the excitatory drive *I*_E_ (A2):

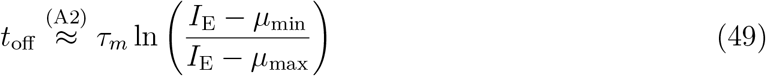

Together with the assumed downstroke duration of Δ we arrive at an analytical estimate for the oscillation period *T* and hence the network frequency *f*_net_ (see Eq. (34)). Overall we have derived analytical expressions for *μ*_max_, *μ*_min_, *t*_off_, *f*_net_ as functions of the external drive *I*_E_, which characterize the oscillatory dynamics. To evaluate the accuracy of our analytical approximation we integrate the DDE numerically (Eq. (33), Fig. 8, green regime of period-1 oscillations) and determine *μ*_max_, *μ*_min_ and *f*_net_. We find that the errors introduced by our simplifying assumptions (A1)-(A3) are small (Fig. 10, dashed lines vs square markers).

**Figure 10:**
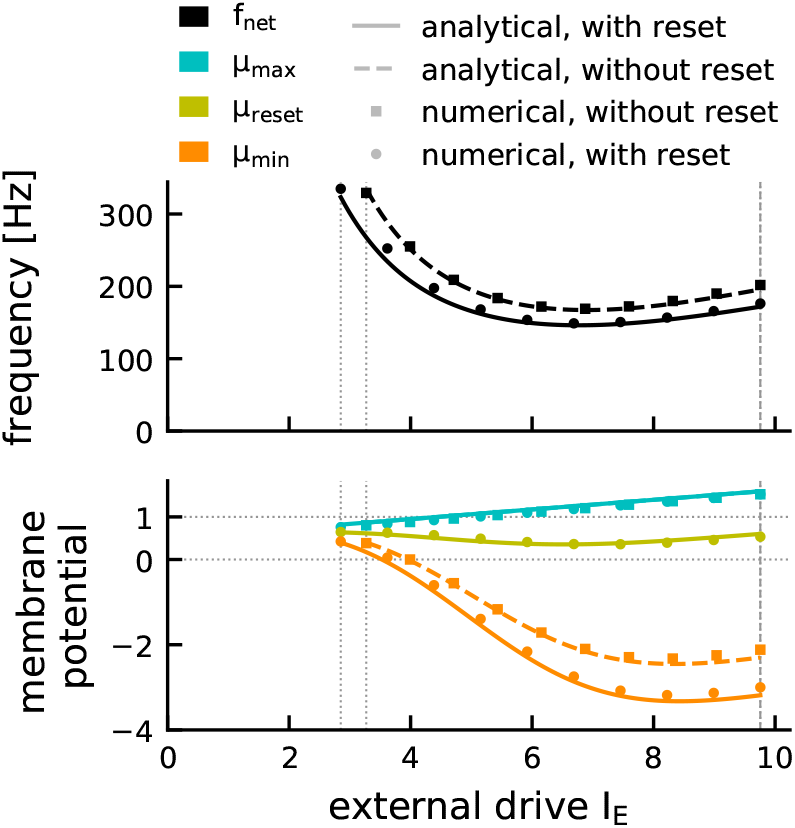
Analytical vs numerical evaluation of oscillatory solutions in the Gaussian-drift approximation. Network frequency (top) and dynamics of the mean membrane potential (bottom) quantified in terms of its periodic local minimum *μ*_min_ (orange) and local maximum *μ*_max_ (cyan) for different levels of external drive *I*_E_. The analytical approximations (solid lines: with reset, dashed lines: without reset) are very close to the results of numerical integration of the DDE Eq. (33) (round markers: with reset, square markers: without reset). Including the reset does not affect *μ*_max_ but leads to a decrease in *μ*_min_ (Eq. (52) vs (45)) and thus a decrease in network frequency. Results are shown in the relevant range of external drives 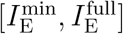 (vertical dotted lines). For parameters see Table 2.

#### Accounting for the reset mechanism

The quantitative accuracy of our Gaussian-drift approximation with respect to the original spiking network can be increased by adding a phenomenological account for the reset on the population level. During the population spike (*t* < *t*_off_) the single unit reset has little influence on the population rate dynamics, since units spike at most once per cycle. At time *t*_off_ the population spike ends and the integral over the suprathreshold portion of the Gaussian density of membrane potentials corresponds to the fraction of units that have spiked (the *saturation*):

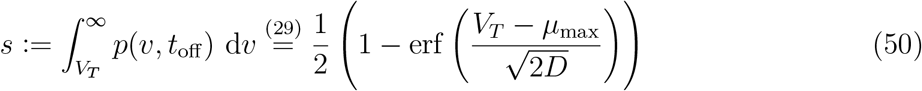

(see Fig. 9C, cyan hatched area). Taking into account the reset mechanism at this point would mean shifting the suprathreshold portion of *p*(*V, t*_off_) downwards by an amount *V_T_*–*V_R_* (Fig. 9C, gray hatched area), essentially splitting the voltage distribution into two pieces, corresponding to silent units (Fig. 9C, non-hatched area under cyan Gauss) and units that have spiked and been reset (Fig. 9C, gray hatched area). To preserve our simplified framework of a unimodal, Gaussian voltage distribution, we will instead assume that the Gaussian voltage distribution is reset as a whole, to a new mean membrane potential *μ*_reset_ given by the average of the two distribution pieces (“silent” and “spike+reset”):

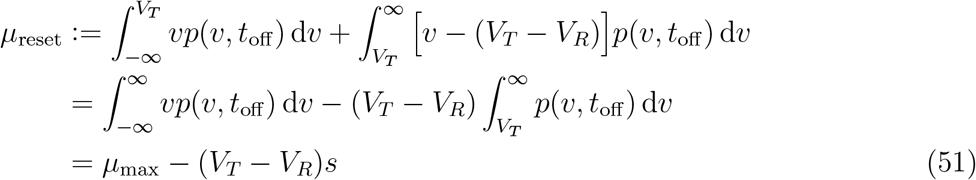

This phenomenological account for the reset contains the implicit assumption that in between population spikes the membrane potential distribution “spends enough time” subthreshold that the bimodality created by the reset mechanism vanishes due to diffusion and the distribution becomes roughly Gaussian again. This assumption is satisfied for a relatively large portion of the parameter space spanned by noise intensity, coupling strength, and reset potential (see Supplementary Information).

The introduction of the population-reset requires an adjustment of the definition of *μ*_min_ (Eq. (42)): Instead of using *μ*_max_ as the initial value when integrating the feedback inhibition during the downstroke, we will now use the reset potential: 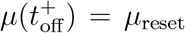. Therefore, *μ*_min_ = *μ*(*t*_off_ + Δ) is given by

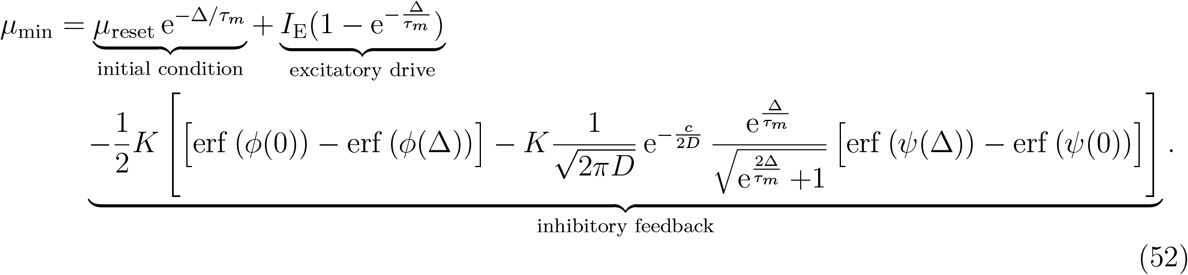

Except for the initial condition term, all other terms remain unchanged (cf. Eq. (45)). Since *μ*_reset_ < *μ*_max_ (Eq. (51)), the introduction of the reset *de*creases our estimate of the local minimum *μ*_min_. This leads to an *in*crease of the upstroke time *t*_off_ required for the mean membrane potential to rise from *μ*_min_ to *μ*_max_ (Eq. (49)), and hence to a *de*crease in the network frequency (Eq. (34), see Fig. 10).

#### The point of full synchrony

Our analytical ansatz allows for a straightforward prediction of the point of full synchrony and its parameter dependencies. As mentioned before, the integral over the suprathreshold-portion of the membrane potential density at the end of the population spike corresponds to the fraction *s* of active units (saturation, Eq. (50)). In the strict sense of

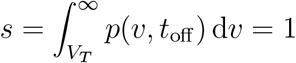

full synchrony can never be reached, since the Gaussian probability density *p* approaches zero only in the limit *v* → ±∞. We can however define *approximate* full synchrony as the state where only the 0.13th percentile of the distribution remains subthreshold:

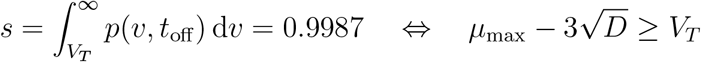

Using our mapping from external drive to *μ*_max_ (Eq. (40)), we can derive a closed-form expression for the external drive that is required to achieve full synchrony:

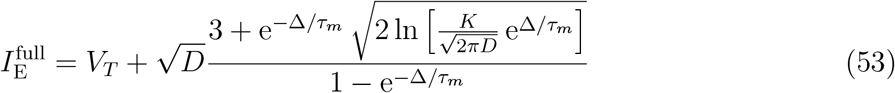

#### Range of applicability of the Gaussian-drift approximation

There are two main constraints on the applicability of our theory:

(a) Since we assume that units spike at most once per cycle, the theory is only valid up to the point of full synchrony 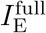 where network frequency and mean unit firing rate coincide.

(b) The assumption of a unimodal distribution of membrane potentials is only valid if, in between population spikes, the bulk of the membrane potential distribution is pushed sufficiently below threshold such that it can diffuse back to approximately Gaussian shape. We will thus require that at its lowest point the Gaussian density is almost entirely subthreshold:

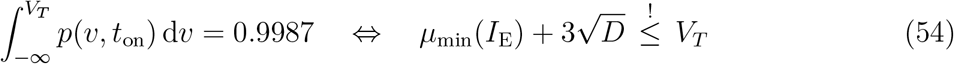

Criteria (a) and (b) yield a finite range 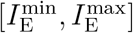 of external drives for which the theory applies. Since for most parameter settings *μ*_min_ is an almost monotonically decaying function of the drive (see Fig. 10 and Results), constraint (b) is usually only relevant for the lower boundary 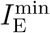 of the drive, while the upper boundary is determined by constraint (a): 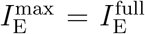 (Fig. 10, see however Fig. S1D, bottom right).

#### Quantifying performance

We quantify the performance of our Gaussian-drift approximation across the range of drives for which

a. the theory applies 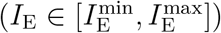
b. the spiking network has not crossed the point of full synchrony 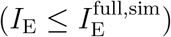

The size of this regime varies for different parameter settings. To ensure comparability we interpolate the results of all spiking network simulations to the same fine resolution:

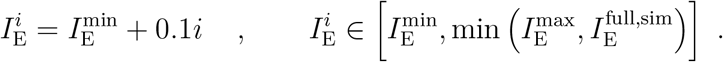

We then compute the average relative error of the estimated network frequencies for each parameter setting:

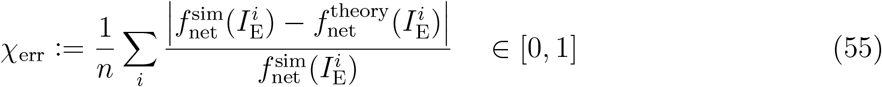

We introduce a second score to quantify what portion of the relevant range of spiking network dynamics (from the Hopf bifurcation, 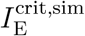, to the point of full synchrony, 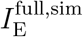) is covered by the theory:

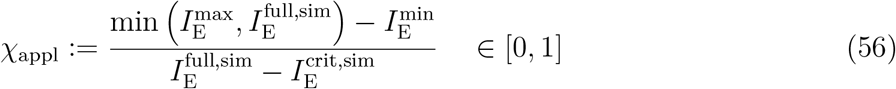

We define an overall performance index as

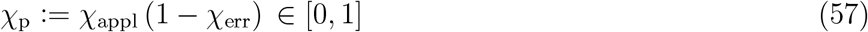

The larger the performance index *χ*_p_ the better the Gaussian-drift approximation captures the spiking network dynamics. A systematic evaluation of the performance of the Gaussian-drift approximation for different network parameters and levels of drive can be found in the Supplementary Material (Fig. S1, Fig. S2).

### Analytical approximation of oscillation dynamics for linear drive

We want to characterize the transient dynamics of any cycle *i* with initial mean membrane potential 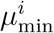 and linear drive 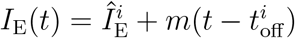 by deriving functions

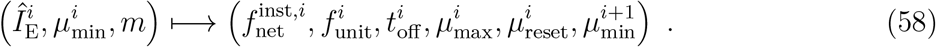

Here 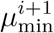 refers to the mean membrane potential reached at the end of cycle *i*, which potentially serves as the initial condition for the next cycle.

#### Step 1

The local maximum of the mean membrane potential 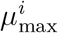 can be found with the same ansatz that was used for constant drive:

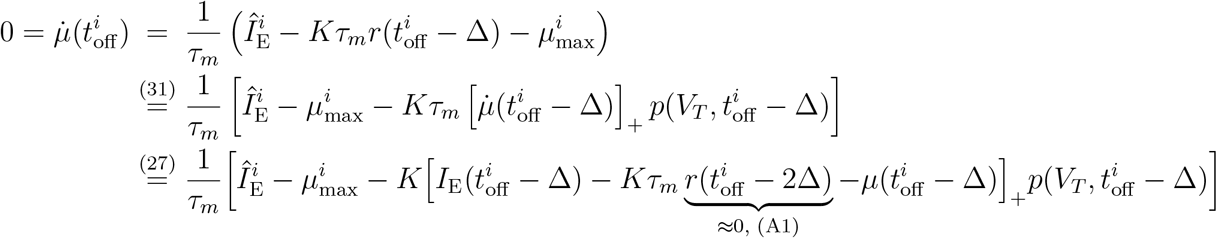

Again we truncate the recurrent expression for the total current two Δ-time windows before the end of the population spike (A1):

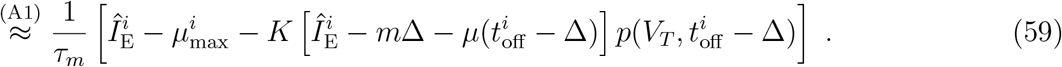

Again we approximate the trajectory of *μ* during the upstroke based on relaxation towards only the excitatory drive, which is now a linear function of time (cf. (A2)). Using the beginning of cycle *i* as the time origin *t* = 0, we obtain

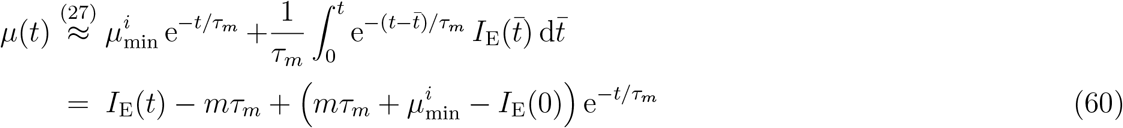

Under this approximation 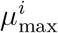 can be written as

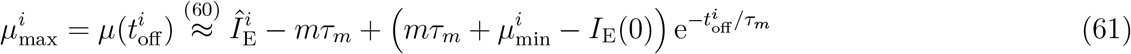

and the trajectory before time 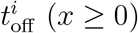 can be approximated as:

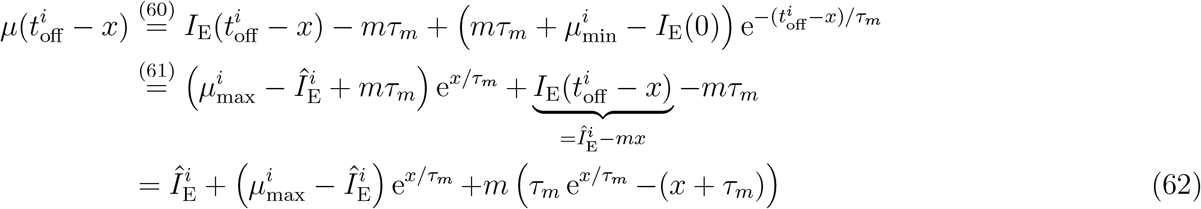

Inserting the above expression in the local maximum condition (Eq. (59)) yields:

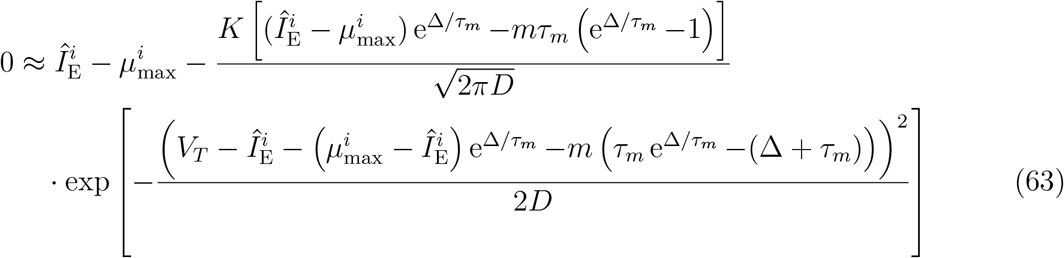

For constant drive (*m* = 0, 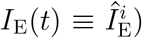) we recover Eq. (39), and thus the asympotic solution 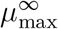. For *m* ≠ 0, we can solve Eq. (63) for small *m*. To this end, we insert the perturbation series 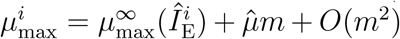 and only keep the terms linear in *m*. Using Eq. (39), we obtain to first order in *m*

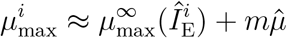

with

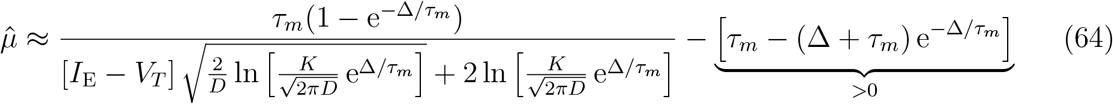

Since the drive is strongly superthreshold 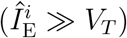 the first order deviation 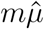 of 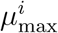 from its asymptotic value 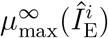 is generally very small. For biologically plausible parameters (*e.g*. a synaptic delay Δ that is not too small), the second term of Eq. (64) is slightly larger than the first. Thus, 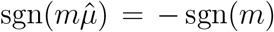, *i.e.* 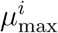 is slightly smaller than its asymptotic value for linearly *in*creasing drive, and slightly larger otherwise (see Fig. 6C, cyan dots vs line).

Since already the zeroth-order approximation 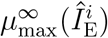 is close to the numerical solution 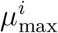 of Eq. (63), and the reset mechanism remains unchanged, this implies that also 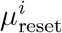 is close to 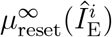.

The duration of the upstroke 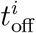 can be obtained from Eq. (61) taking into account that 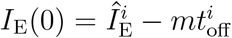:

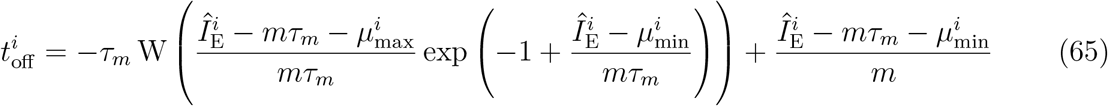

where W is the Lambert W function, which has solutions for arguments > − exp(−1). For positive slope *m* > 0 and low drive 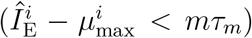 this introduces a constraint on the initial value:

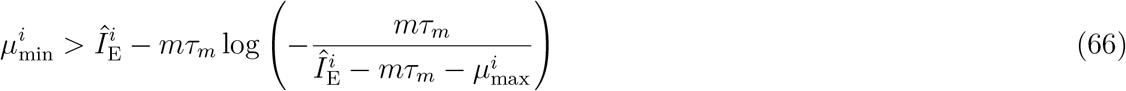

(cf. Fig. 6B, 7Ai).

The instantaneous frequency of the cycle follows as

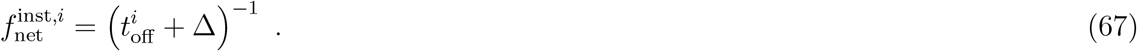

Note that for linearly changing drive, the asymptotic initial value 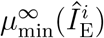 no longer implies the asymptotic frequency 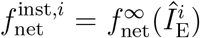. If the drive increases linearly with a positive slope *m* > 0, the correct initial value 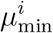 leading to 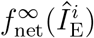 (Fig. 6B, left, white line) is larger than 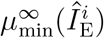 (Fig. 6B, left, black line). Conversely, if the drive decreases linearly (*m* < 0), the correct initial value 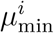 leading to 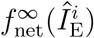 is smaller than 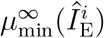 (Fig. 6B, right, white vs black line). The correction of the initial condition can be understood as follows: if *e.g*. the slope is positive, *m* > 0, the driving current *I_E_*(*t*) is smaller than 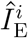 during the upstroke of *μ*(*t*), which lasts until time 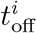 (cf. Eq. (12) and Fig. 6A, left). A smaller driving current results in a slower increase of *μ*(*t*), (cf. Eq. (8a)), and hence a longer rise time from 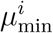 to 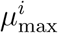. In order to match the asymptotic frequency, the initial value must therefore be chosen larger than 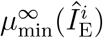 so as to compensate the slower increase of *μ*(*t*) during the rising phase. A similar argument holds when the driving current is decreasing (*m* < 0).

#### Step 2

We will now compute the mean membrane potential 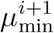 that is reached at the end of a cycle *i*. Note that this step is technically not necessary to understand the instantaneous frequency 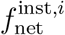 of an isolated cycle *i*, which we already derived above as a function of the reference drive 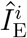, the slope *m*, and an arbitrary initial condition 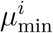 (Eq. (67)). It is however of interest to demonstrate that the mean membrane potential 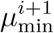 at the end of the cycle is close to the asymptotic reference 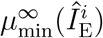, since it serves as the initial condition of the next cycle. It is this property, together with the dependence of 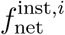 on the initial condition 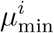, that implies the emergence of IFA under a piecewise linear drive that first increases, and then decreases over multiple consecutive ripple cycles (cf. Results, Fig. 6C).

We find 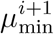 by integrating the total current during the downstroke:

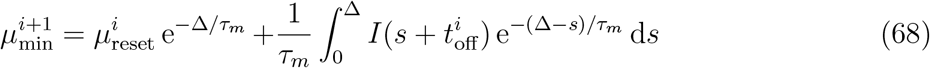

The total current *I* can be split into two parts: *I*^stat^, which is approximately equal to the feedback current for constant drive 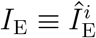 (Eq. (43), except for slight deviations in 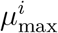), and an additive new term *I^m^* caused by the linear change in the external drive:

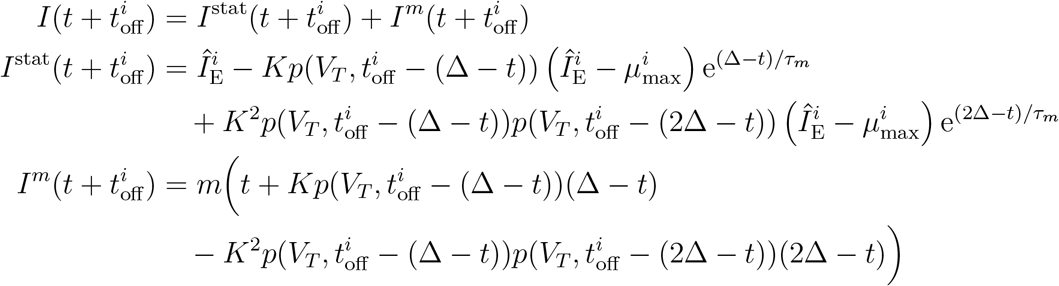

For constant drive (*m* = 0) we recover the asymptotic solution 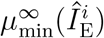 (Eq. (52)). For *m* ≠ 0 we integrate Eq. (68) numerically and find that 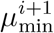 is indeed close to the asymptotic solution 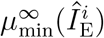. One can infer intuitively, that the sign of the (small) deviation of 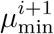 from 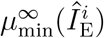 equals the sign of the slope *m*: If the drive *in*creases linearly, the drive during the upstroke of the mean membrane potential 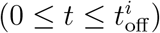 is *below* the reference drive 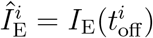 (Eq. (12)). Thus *μ* rises towards a slightly smaller maximum 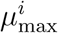 (Eq. (64)) and with slightly reduced speed, which decreases the resulting population rate and thus the inhibitory feedback (Eq. (31), Eq. (22b)). Furthermore, the excitatory drive during the feedback-induced downstroke of *μ*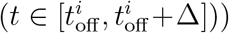 is stronger than the reference drive (Eq. (12)). It follows that the downstroke of the mean membrane potential is reduced, and thus 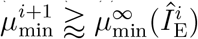. The opposite argument can be made for linearly *de*creasing drive.

At this point it is clear that a piecewise linear drive, that first increases and then decreases over multiple ripple cycles (Eq. (17)) will inevitably induce IFA. It is also clear, that this IFA asymmetry will vanish in the limit of infinitely slow drive (|*m*| → 0).

Showing a concrete example of IFA requires a forward integration of the network response over multiple cycles. Note that our ansatz centered around the reference point 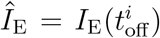 (Eq. (58)) does not allow a straight-forward calculation of such trajectories over *multiple, consecutive* cycles. For any *individual* cycle *i* we can understand the frequency difference 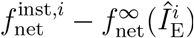 and how it depends on the drive 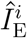, its slope *m* and the initial condition 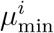. We can also infer the drive at the beginning and end of the cycle as 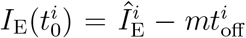 and 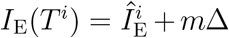. In retrospect, we could match cycles self-consistently such that the drive at the end of one cycle equals the drive at the beginning of the next: 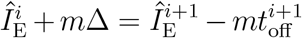. Instead we chose to compute the example trajectory shown in Fig. 6C numerically by integrating the DDE (Eq. (33)), enforcing the population reset (Eq. (51)) at the end of each population spike.

#### Comparing theory and simulation for piecewise linear drive

Since the theory provides a discrete, cycle-wise estimate of the instantaneous network frequency, we compare the result to the discrete estimate of instantaneous frequency in the simulations, based on the inverse of the distances between consecutive peaks in the oscillatory population rate. In the spiking network, SPW-like drive is modeled as a piecewise linear double-ramp with an intermediate plateau phase of 20 ms (Eq. (17)). The Gaussian-drift approximation is used to estimate the instantaneous network frequencies separately for the rising and falling phases of the drive (linear increase or decrease with slope ±*m*). The plateau phase is ignored, since the network frequencies rapidly converge to the asymptotic frequency associated to the drive during the plateau phase. In both simulation and theory IFA is quantified by computing a linear regression slope over the instantaneous frequencies. The theoretically estimated instantaneous frequencies are shifted in time to account for a hypothetical plateau phase of 20 ms in between up- and downstroke and allow full comparability with the simulation results.

For every theoretical instantaneous frequency estimate 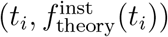 an error is calculated relative to the average instantaneous frequencies observed in the spiking simulation around the same time point (*t_i_* ± 1.5 ms):

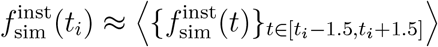

The average relative error of the theoretical estimate, compared to the simulations, is then computed as

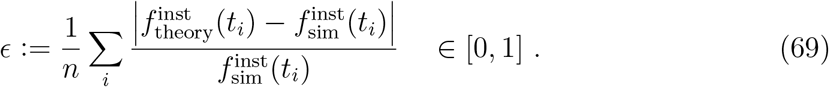

(see Table 1).

## Acknowledgements

We thank José R. Donoso, Naomi Auer and Gaspar Cano for critical discussions and editing suggestions.

## Supplementary

### Performance evaluation of the drift-based approximation

We confirmed with numerical simulations that for strong enough drive the Gaussian-drift approximation works for a wide parameter regime. Our spiking network model has only four parameters: the noise intensity *D*, the inhibitory coupling strength *K*, the synaptic delay Δ, and the reset potential *V_R_* (see Eq.(3)). Since time can be rescaled to units of the membrane time constant, we do not count the membrane time constant as an independent parameter. We show here a two-dimensional parameter exploration covarying the noise intensity *D* and the inhibitory coupling strength *K* (Fig. S1). Subsequently we will briefly comment on the role of the synaptic delay and the reset potential.

To quantify the performance of the Gaussian-drift approximation, we introduce a performance index that takes into account the error in the estimate of the network frequency and the proportion of the relevant range of external drives (from the Hopf bifurcation up to the point of full synchrony) that is covered by the theory (Methods, Eq. (57)).

We find that performance is good for a wide range of parameters (Fig. S1B). As expected from a drift-based approximation, the performance decreases for larger noise intensity (Fig S1, large 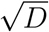). At high noise and weak coupling the range of external drives, for which the theory applies, decreases markedly. The wider the Gaussian density and the weaker the inhibitory feedback, the harder it is to satisfy our requirement (b) of the bulk of the membrane potential density being pushed subthreshold in between population spikes (Fig. S1D). In the extreme case of 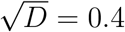, *K* = 2 this criterion is never fulfilled (Fig. S1B-D, bottom right).

We also observe a dip in performance for low noise and low coupling strength (Fig. S1B-D, bottom left). This, however, does not reflect a shortcoming of our approximation but rather a departure of the spiking network dynamics from the regime of interest: If the noise is very low relative to the distance between threshold and reset, the portion of active units in a given cycle is reset far below the portion of silent units, which remain just below threshold. If in addition the synaptic coupling is also weak, these disjoint pieces will not merge back together before the previously silent portion hits the threshold and starts the next population spike. This leads to a membrane potential distribution that is multimodal at all times and where units that have spiked in one cycle are less likely to spike in the next. Such a *clustered* activity (see also Brunel and Hansel, 2006) does not correspond to our regime of interest and can of course not be captured by our approximation. To get back to the regime of interest for the given level of low noise and low coupling, one could increase the reset potential, essentially introducing a third dimension to our parameter exploration.

Our parameter exploration reveals that the network frequency beyond the Hopf bifurcation depends strongly on the inhibitory coupling strength (Fig. S1C, variation along vertical axis *K*). Previous studies focusing on linear stability analysis around the Hopf bifurcation (Brunel and Hakim, 1999; Brunel and Wang, 2003) have suggested that the network frequency is set primarily by the (fast) synaptic time constants and depends only weakly on other parameters, which our simulations confirm (see Fig.S1C, red markers). Here we see, however, that further away from the bifurcation other parameter dependencies *do* play a role. For strong coupling (*K* = 50, Fig. S1C, top row) we even observe a drop in the network frequencies to slow gamma range. Our approximation captures this dependency very well.

**Figure S1:**
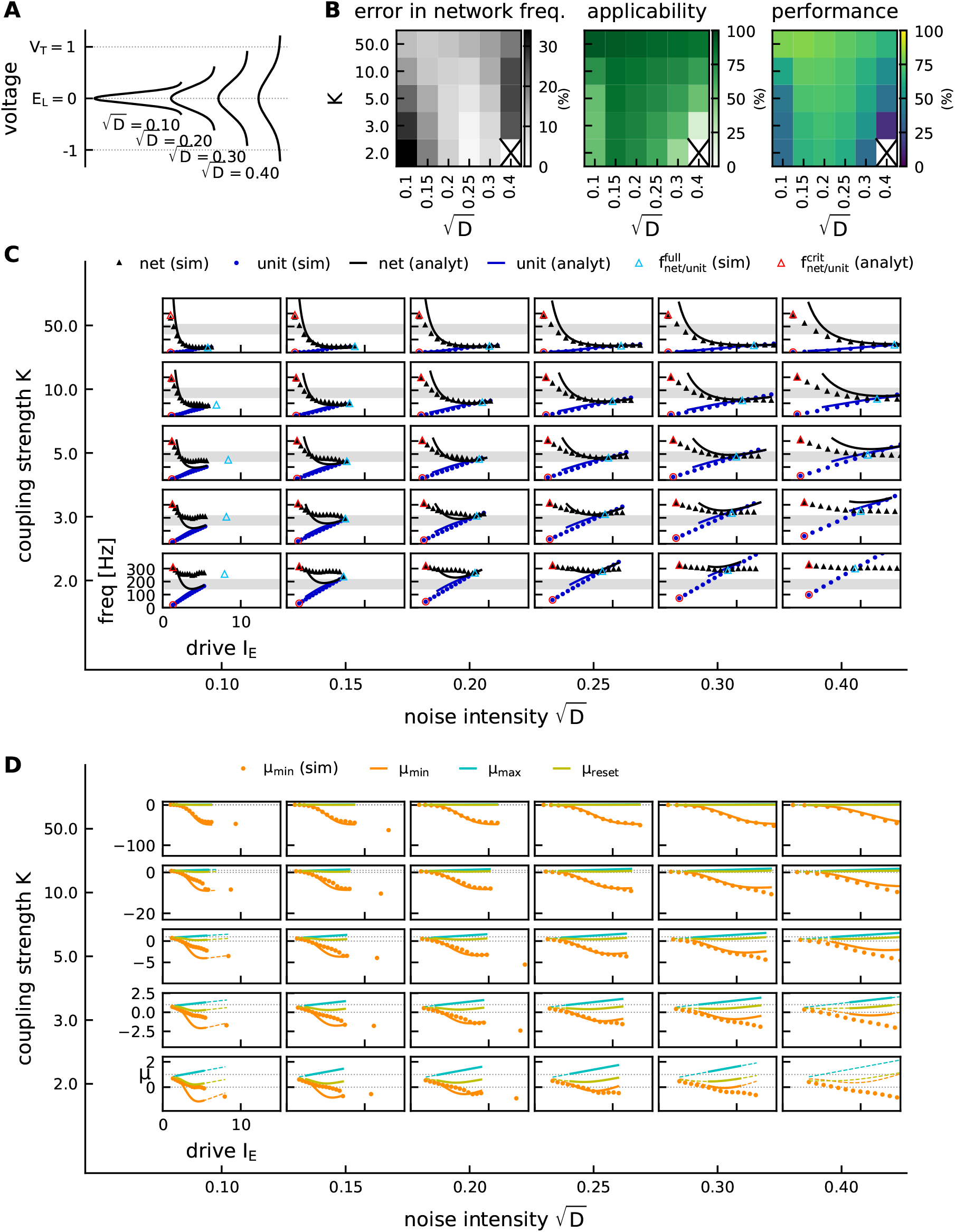
Performance of the Gaussian-drift approximation in a 2D parameter exploration. Noise intensity *D* and coupling strength *K* were covaried in the range 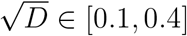 and *K* ∈ [2, 50]. **A**, Visualization of the width of the Gaussian voltage density *p*(*V, t*) for different noise intensities *D*. **B**, performance index reveals optimal parameter regime in terms of approximation error and applicability of the theory. **C**, comparison of network frequencies (black) and unit firing rates (blue) in theory (line) and simulation (markers). Red markers: Hopf bifurcation. Blue triangle: point of full synchrony in spiking network simulation. All theory curves are shown for the respective range 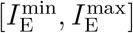 (see Methods, Eq. (54)). **D**, local extrema *μ*_max_, *μ*_min_ and reset *μ*_reset_ of the mean membrane potential. Dashed colored lines: theory does not apply. Full colored lines: theory applies, 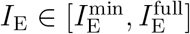. Note that *μ*_min_ is a quasi monotonically decaying function of the drive, except for very weak coupling.

The point of full synchrony is predicted well (Fig. S1C). Its parameter dependencies are illustrated in Fig. S2: while *quantitatively* the theoretical estimate produces an error that becomes larger for stronger noise (Fig. S2, *error*), the *qualitative* dependency of the point of full synchrony on the noise and coupling strength is captured well (Fig. S2, *theory* vs *simulation*). The theory predicts that stronger external drive is required to achieve full synchrony, if the noise is stronger, which is confirmed in simulations (Fig. S2 bottom). A similar, albeit weaker, dependence can be found for the inhibitory coupling strength: If the network is coupled more strongly, stronger external drive is required to achieve full synchrony.

**Figure S2:**
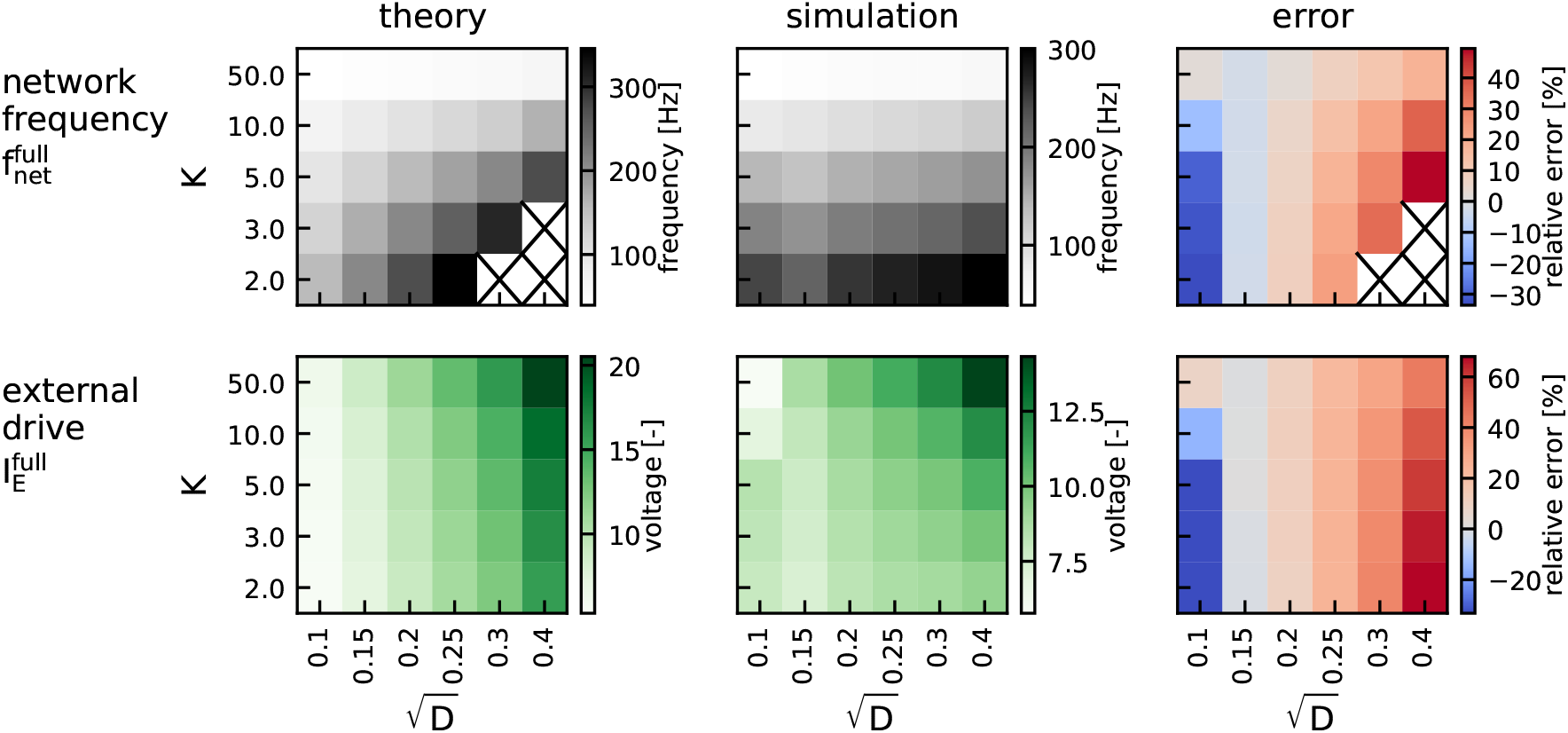
Parameter dependencies of the point of full synchrony. External drive 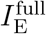 at which full synchrony is reached (bottom) and the corresponding network frequency 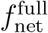 (top), as predicted by theory (left, Eq.(53)) vs. as observed in spiking network simulation (middle). The right panel shows the relative deviation of the theoretical prediction from the simulation result. Same parameter exploration as in Fig.S1. Crosses mark the parameter settings for which the point of full synchrony lies outside the regime of applicability of our theory 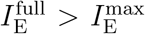, cf. Fig. S1D).

What about the remaining two parameters, Δ and *V_R_*? We have covaried the synaptic delay in a biologically plausible range (Δ ∈ [0.5, 2] ms) with either the noise intensity or the coupling strength (not shown here). The performance of the Gaussian-drift approximation is largely unaffected by the synaptic delay, which merely shifts the overall network frequencies to higher or lower values as predicted by Brunel and Hakim (1999). As mentioned before, the reset potential *V_R_* can introduce permanent multimodality in the distribution of membrane potentials if it is far from threshold relative to noise intensity and coupling strength. As long as approximate unimodality is ensured, the reset potential does not influence the performance of the approximation much.

