## Supplementary Material for "Intra-ripple frequency accommodation in an inhibitory network model for hippocampal ripple oscillations"

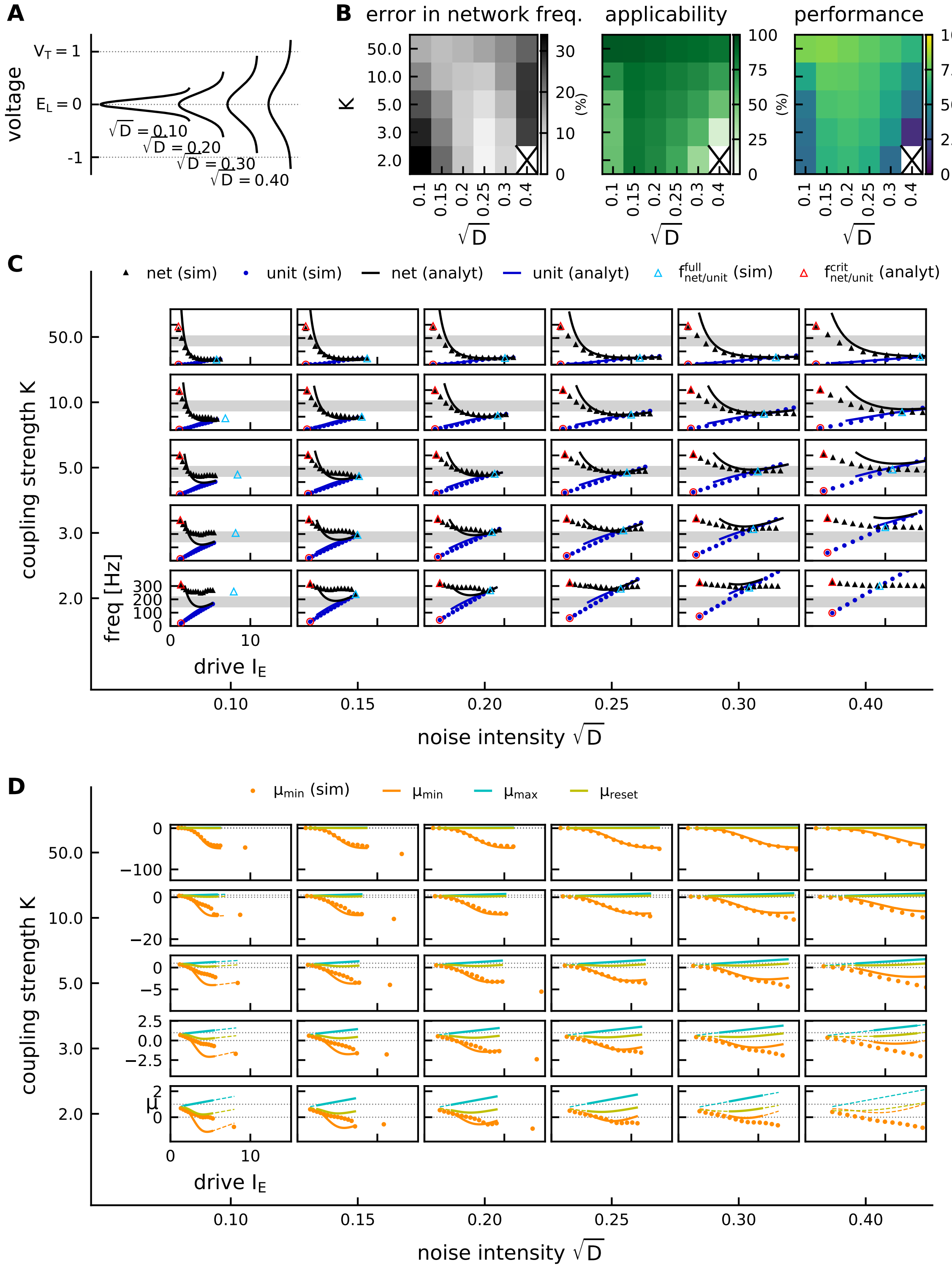


**Figure S1: Performance of the Gaussian-drift approximation in a 2D parameter exploration.** Noise intensity $D$ and coupling strength $K$ were covaried in the range $\sqrt{D} \in [0.1, 0.4]$ and $K \in [2, 50]$. **A,** Visualization of the width of the Gaussian voltage density $p(V, t)$ for different noise intensities $D$. **B,** performance index reveals optimal parameter regime in terms of approximation error and applicability of the theory. **C,** comparison of network frequencies (black) and unit firing rates (blue) in theory (line) and simulation (markers). Red markers: Hopf bifurcation. Blue triangle: point of full synchrony in spiking network simulation. All theory curves are shown for the respective range $[I_E^{min}, I_E^{max}]$ (see Methods, Eq. (54)). **D,** local extrema $\mu_{max}, \mu_{min}$ and reset $\mu_{reset}$ of the mean membrane potential. Dashed colored lines: theory does not apply. Full colored lines: theory applies, $I_E \in [I_E^{min}, I_E^{full}]$. Note that $\mu_{min}$ is a quasi monotonically decaying function of the drive, except for very weak coupling.

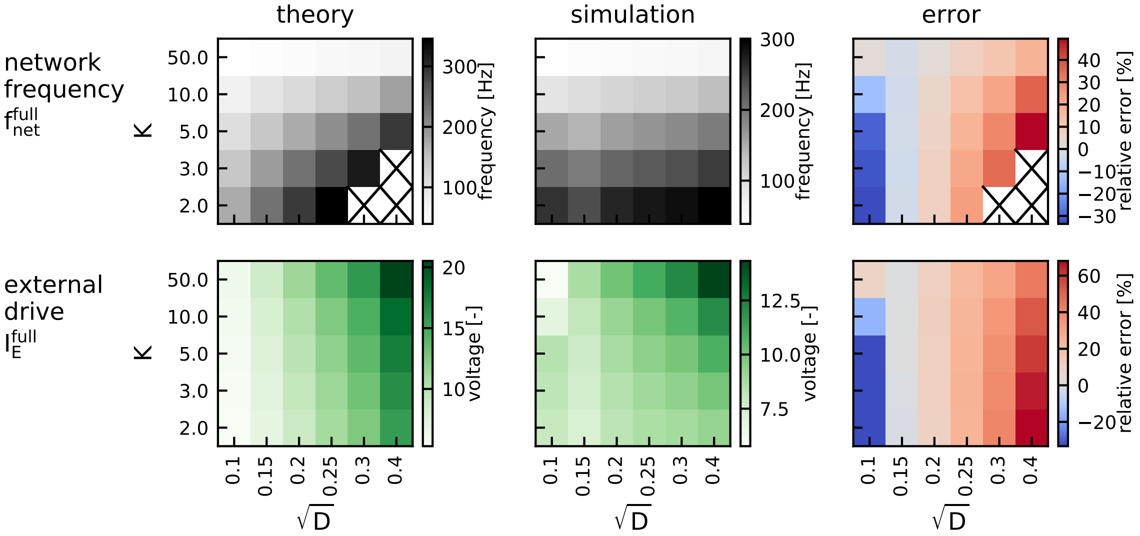


**Figure S2: Parameter dependencies of the point of full synchrony.** External drive $I_E^{full}$ at which full synchrony is reached (bottom) and the corresponding network frequency $f_{net}^{full}$ (top), as predicted by theory (left, Eq. (53)) vs. as observed in spiking network simulation (middle). The right panel shows the relative deviation of the theoretical prediction from the simulation result. Same parameter exploration as in Fig.S1. Crosses mark the parameter settings for which the point of full synchrony lies outside the regime of applicability of our theory ($I_E^{full} > I_E^{max}$, cf. Fig. S1D).
